# Increasing the throughput of sensitive proteomics by plexDIA

**DOI:** 10.1101/2021.11.03.467007

**Authors:** Jason Derks, Andrew Leduc, Georg Wallmann, R. Gray Huffman, Matthew Willetts, Saad Khan, Harrison Specht, Markus Ralser, Vadim Demichev, Nikolai Slavov

## Abstract

Current mass-spectrometry methods enable high-throughput proteomics of large sample amounts, but proteomics of low sample amounts remains limited in depth and throughput. To increase the throughput of sensitive proteomics, we developed an experimental and computational framework, plexDIA, for simultaneously multiplexing the analysis of both peptides and samples. Multiplexed analysis with plexDIA increases throughput multiplicatively with the number of labels without reducing proteome coverage or quantitative accuracy. By using 3-plex nonisobaric mass tags, plexDIA enables quantifying 3-fold more protein ratios among nanogram-level samples. Using 1 hour active gradients and first-generation Q Exactive, plexDIA quantified about 8,000 proteins in each sample of labeled 3-plex sets. plexDIA also increases data completeness, reducing missing data over 2-fold across samples. We applied plexDIA to quantify proteome dynamics during the cell division cycle in cells isolated based on their DNA content; plexDIA detected many classical cell cycle proteins and discovered new ones. When applied to single human cells, plexDIA quantified about 1,000 proteins per cell and achieved 98 % data completeness within a plexDIA set while using about 5 min of active chromatography per cell. These results establish a general framework for increasing the throughput of sensitive and quantitative protein analysis.

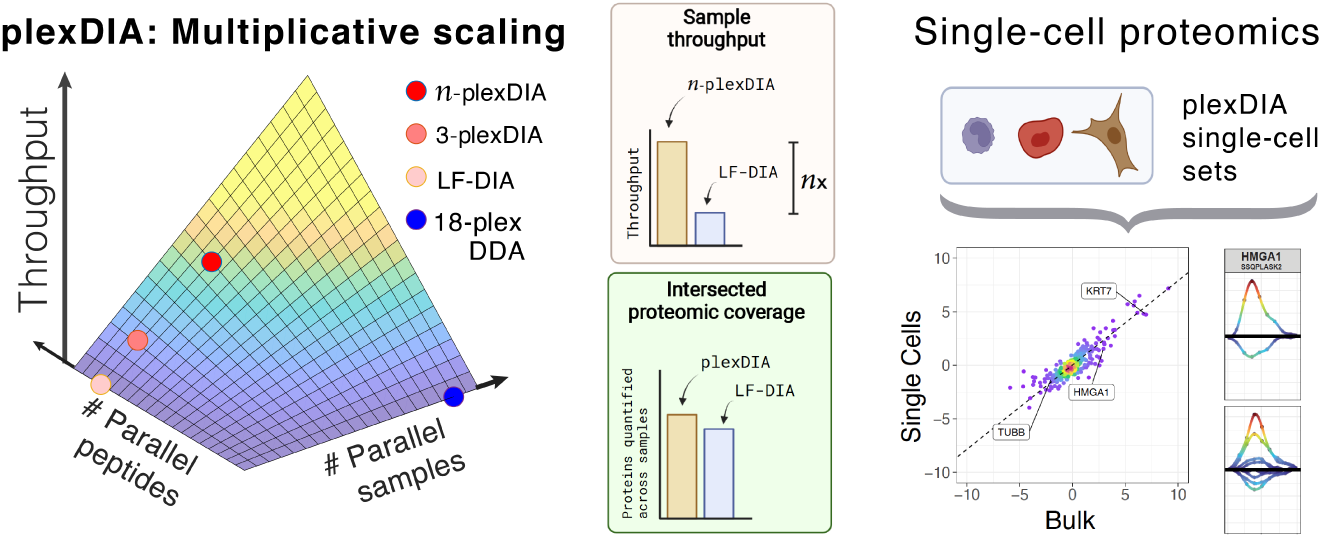

## Introduction

Mass-spectrometry (MS) methods can achieve deep proteome coverage^1, 2^, low missing data^3^, high throughput^4, 5^, and high sensitivity^6^. However, simultaneously achieving all these objectives is an outstanding challenge^7, 8^. Resolving this challenge will empower biomedical projects that are impractical with current methods^8^, especially those that require single-cell protein analysis^9–11^. Towards this goal, the throughput of sensitive protein analysis may be increased by different strategies: (i) increasing sample throughput and robustness by chemical labeling, and (ii) decreasing MS analysis time per sample by simultaneous (parallel) analysis of multiple peptides. These strategies are complementary, and we sought to combine them to achieve a multiplicative increase in the rate of quantifying the proteomes of limited sample amounts.

Chemical labeling is often used with data-dependent acquisition (DDA) to increase throughput via parallel sample analysis (Fig. 1a) and to control for shared artifacts, such as disturbances in peptide separation and ionization^12–14^. Since quantifying a mammalian proteome requires analyzing hundreds of thousands of precursor ions and DDA methods analyze one precursor per MS2 scan, even the most optimized DDA methods require up to a day of LC-MS/MS for deep proteome analysis^1^. Nonisobaric labels, such as mTRAQ and dimethyl labeling allow for sample multiplexing but further increase the number of precursor ions and thus the time needed for MS1- multiplexed DDA analysis^15^. In contrast, approaches using isobaric labels (such as TMT; tandem mass tags) do not increase the number of distinguishable precursor ions and can reduce the analysis time per sample^16, 17^, albeit quantification with TMT is often significantly affected by coisolation interference^13, 18^.

**Figure 1.**
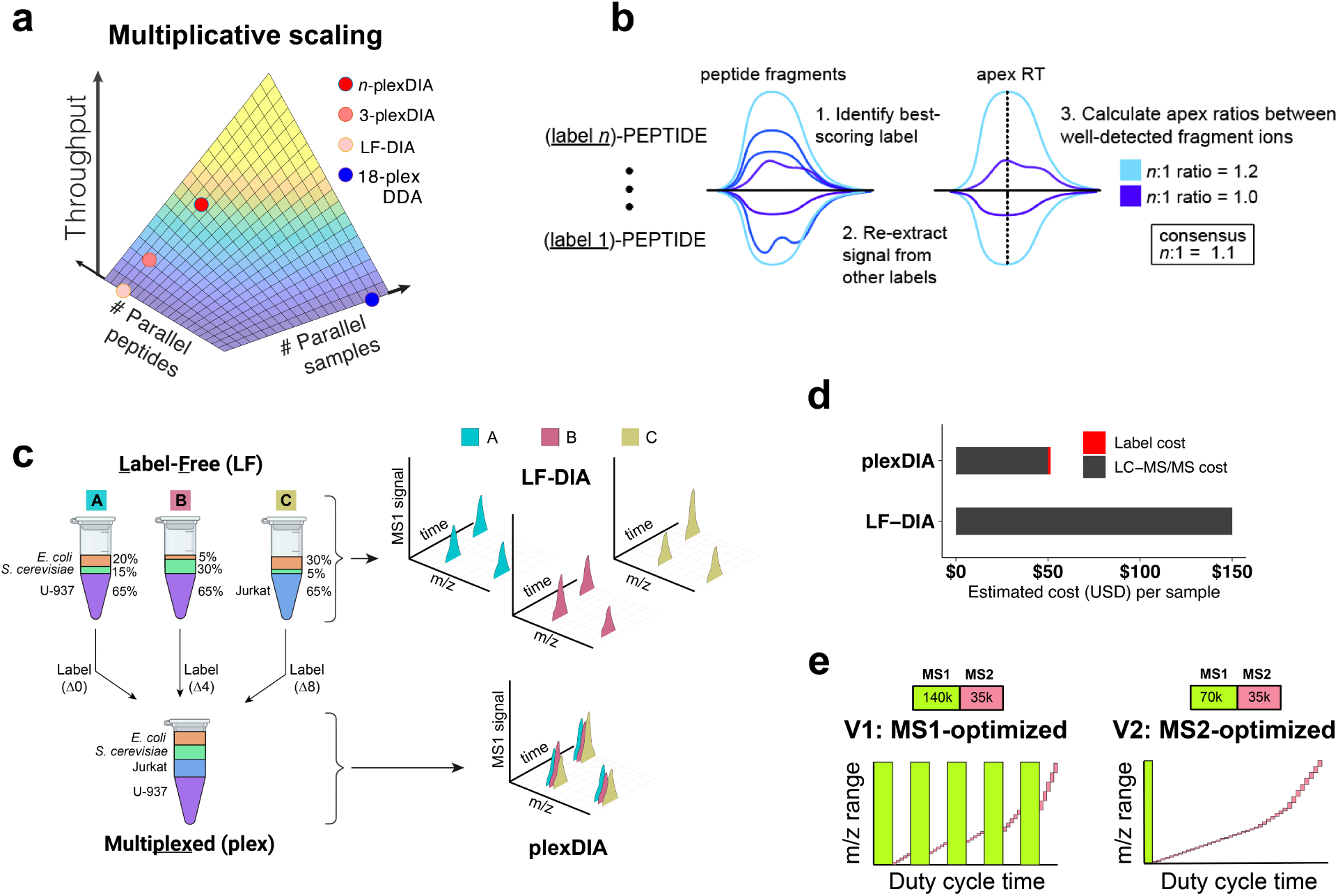
Experimental design for acquisition and evaluation of plexDIA data. (**a**) The throughput of MS proteomics can be increased by parallel analysis of multiple peptides or by parallel analysis of multiple samples. plexDIA aims to combine both approaches to achieve multiplicative gains. (**b**) Precursor identifications from one label can be confidently transferred to isotopologous precursors with FDR control. The abundance of labeled precursors can be estimated by the consensus fold-change relative to best quantified isotopologous precursor. (**c**) Standards used for benchmarking LF-DIA and plexDIA quantification were prepared by mixing the proteomes of different species and cell types as shown. LF-DIA analyzed 500ng from each of the 3 samples (A, B, C) separately, while plexDIA analyzed a mixture of these samples labeled with nonisobaric mass tags (mTRAQ). (**d**) Analyzing samples by plexDIA is cheaper than analysis by LF-DIA because running *n* samples in parallel reduces the LC-MS/MS time per sample *n*-fold and the cost of labeling is low. This estimate is based on a facility fee of 150 USD / hour of active gradient. (**e**) We benchmarked the performance of plexDIA with two DIA methods, V1 and V2. V1 is an MS1-optimized method that utilizes frequent, high resolution MS1 scans to facilitate accurate quantification while V2 is an MS2-optimized method which takes a single MS1 scan and more MS2 scans per duty cycle.

The throughput of DDA analysis can be increased by decreasing the ion accumulation times for MS2 scans, though this results in accumulating fewer ions and thus limits sensitivity^7^. Indeed, sensitive analysis of small sample amounts requires (and is thus limited) by long ion accumulation times, which are typically substantially longer than the detection time required by MS detectors^6, 19, 20^. Even with short ion accumulation times for unlimited sample amounts, the requirement to serially analyze hundreds of thousands of precursor ions remains a major challenge for simultaneously achieving high throughput and deep proteome coverage by serial precursor analysis.

A fundamental solution to this challenge is the concept of isolating and analyzing multiple precursor ions simultaneously by data-independent acquisition (DIA)^21^. This concept has been implemented into powerful methods for label-free DIA (LF-DIA) protein analysis^22–26^. Such parallel analysis of peptides decreases the time needed to analyze thousands of precursor ions and makes the throughput of optimized LF-DIA and TMT-DDA workflows comparable (Fig. 1a), allowing routine quantification of about 6,000 proteins in 2 hours^17^. Recent DIA technologies further enabled quantification of over 8,000 proteins in 1.5 hours^27^ and TMTpro tags increased multiplexing to 18-plex for DDA methods^4^. Thus multiplexed DDA and LF-DIA afford comparable throughput, Fig. 1a.

We sought to further increase the throughput of sensitive DIA by multiplexing samples labeled with nonisobaric isotopologous mass tags, capitalizing on the fact that increasing the number of precursor ions does not increase the time needed to analyze them via tandem DIA-MS, in contrast to DDA-MS^15, 21^. This creates a hypothetical possibility that we sought to test: The number of proteins accurately quantified by multiplexed DIA may increase multiplicatively with the number of labels used, Fig. 1a. If feasible, this possibility may enable higher throughput and more sensitive multiplexed proteomics, including single-cell proteomics as previously suggested^7, 28^. While the feasibility of DIA multiplexed by SILAC^29^ or pulsed SILAC^30, 31^ has been clearly demonstrated, its ability to multiplicatively increase quantitative data points remains unclear. Similarly, clever strategies have used both isobaric and isotopologous tags to multiplex DIA, but they have afforded the quantification of relatively few proteins^32–34^. Thus, the potential of multiplexed DIA to increase sample throughput while preserving proteome coverage and quantification accuracy has not been realized due to the increased complexity of DIA data from labeled samples^33–37^.

We hypothesized that an optimized experimental and analytical framework may enable *n*-fold multiplexed DIA to increase *n*-fold the number of accurate protein data points, Fig. 1a. We test this hypothesis for *n* = 3 using amine-reactive nonisobaric isotopologous mass tags (mTRAQ), hoping that this particular choice of mass tags can establish a framework that will in the future generalize to a variety of isotopologous non-isobaric mass tags with higher capacity for multiplexing. Specifically, we sought to develop a general framework and an analysis pipeline to increase the throughput of sensitive and quantitative protein analysis via plexDIA.

## Results

To enhance MS data interpretation, the plexDIA module of DIA-NN capitalizes on the expected regular patterns in the data, such as identical retention times and known mass-shifts between the same peptide labelled with different isotopologous mass tags, Fig. 1b, Fig. S1^7^. DIA-NN uses neural networks to confidently identify labeled peptides, and these identifications are then used to re-extract data for the same peptide labeled with a different tag. Neural networks then calculate false discovery rates for all peptides based on a decoy channel strategy, which is empirically validated by two-species spiked experiment shown in Fig. S2. Despite the *n*-fold increased spectral complexity, the plexDIA framework aims to accurately quantify peptides by calculating ratios of fragments from the most confident isotopologous precursor to the translated isotopologous precursors at the apex where the signal is greatest and the impact of interference is lowest. The mean fragment ratio is used to scale the precursor quantity of the best isotopologous precursor to the less-confident isotopologous precursors, Fig. 1b.

### plexDIA benchmarks

We sought to evaluate whether plexDIA can multiplicatively increase the number of quantitative data points relative to matched label-free DIA (LF-DIA) analysis while maintaining comparable quantitative accuracy. Towards that goal, we mixed proteomes in precisely specified ratios shown in Fig. 1c, thus creating a benchmark of known protein ratios for thousands of proteins spanning a wide dynamic range of abundances, similar to previous benchmarks^23^. Specifically, we made three samples (A, B, and C), each with an exactly specified amount of *E. coli*, *S. cerevisiae*, and *H. sapiens* (U-937 and Jurkat) cell lysate, Fig. 1c. A distinct aspect of this design is the incorporation of human proteomes of different cell types, which affords additional benchmarking for the reproducibility of protein identification across diverse samples and for relative protein quantification.

Each sample was either analyzed by label-free DIA (LF-DIA) or labeled with one of three amine-reactive isotopologous chemical labels (mTRAQ: Δ0, Δ4, or Δ8), Fig. 1c. With this experimental design, plexDIA enables 3-fold reduction in LC-MS/MS time per sample, which provides nearly 3-fold reduction in the overall cost per sample because most of the cost stems from LCMS/MS fees while the cost of labeling is low, Fig. 1d. The combined labelled samples were analyzed by plexDIA, and the result was used to benchmark proteomic coverage, quantitative accuracy, precision, and repeatability across runs relative to LF-DIA of the same samples. LF-DIA and plexDIA were evaluated with two data acquisition methods, V1 and V2, shown in Fig. 1e. V1 included multiple high-resolution MS1 survey scans to increase the temporal resolution of precursor sampling as previously reported^3^ while V2 included more MS2 scans to increase proteome coverage, Fig. 1e; The only difference between the duty cycles of LF-DIA and plexDIA was a 100 m/z increment in the MS1 and MS2 windows of plexDIA to account for the mass of mTRAQ added to the peptides; see methods.

### plexDIA increases throughput multiplicatively

To directly benchmark the analysis of 500 ng protein samples by plexDIA relative to LF-DIA, the multiplexed and label-free samples described in Fig. 1c were analyzed in triplicate by LCMS/MS on Thermo Q-Exactive (first generation) with a 60-min active nano-LC gradient. The throughput increases for duty cycles V1 (Fig. 2) and V2 (Fig. S3) were similar, except that V2 achieved greater proteome coverage with both plexDIA and LF-DIA. The parallel data acquisition by all DIA methods resulted in a greater number of identified peptides and proteins compared to the DDA runs, Fig. 2a,b. Both V1 and V2 resulted in approximately 2.5-fold more precursors and protein data points for plexDIA compared to LF-DIA per unit time, Fig. 2a,b & Fig. S3a,b.

**Figure 2.**
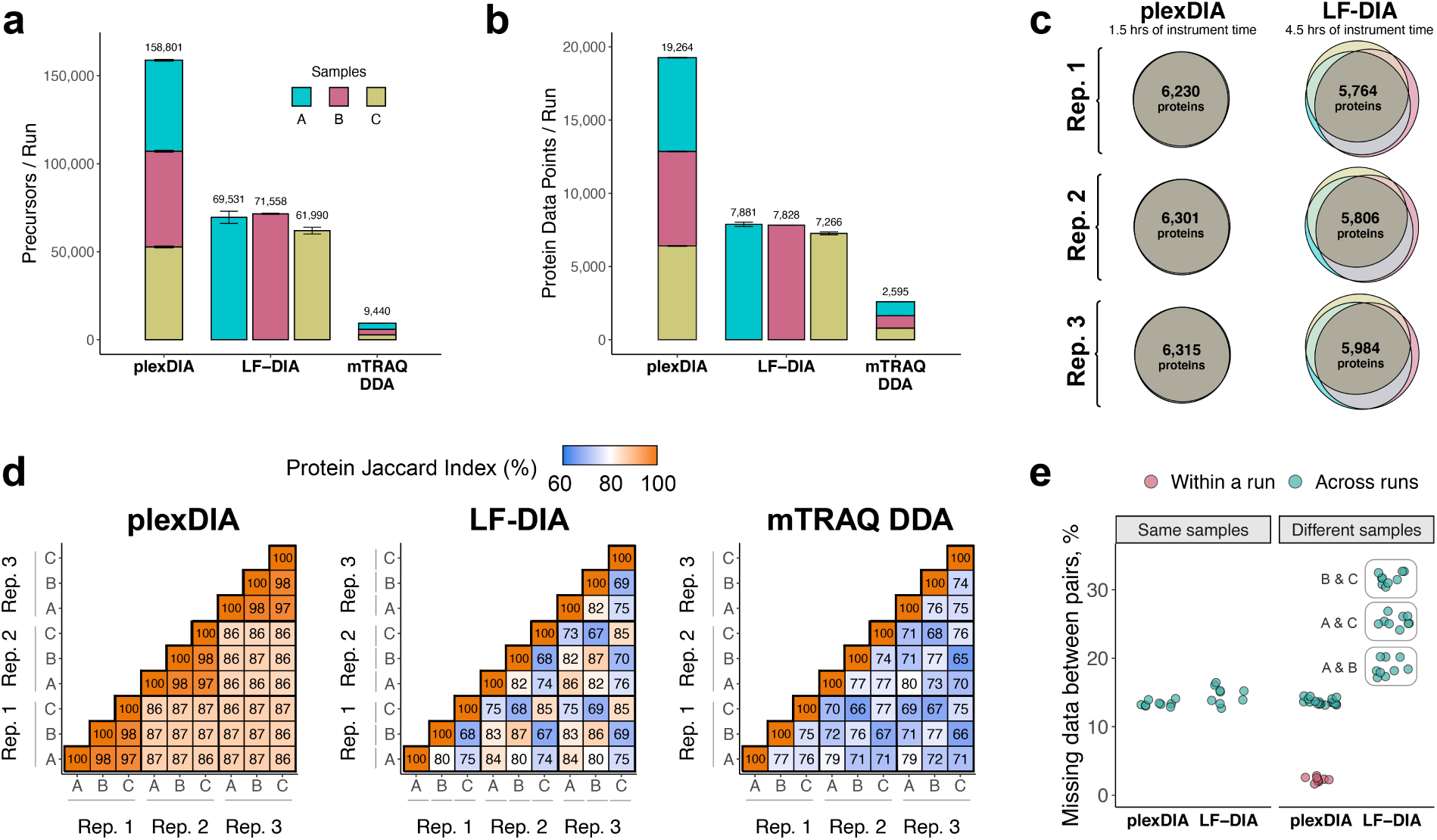
plexDIA proteome coverage and overlap between samples and runs. (**a**) Number of distinct precursors identified from 60 min active gradient runs of plexDIA, LF-DIA, and mTRAQ DDA at 1 % FDR. The DIA analysis employed the V1 duty cycle shown in Fig. 1c. Each sample was analyzed in triplicate and the results displayed as mean; error bars correspond to standard error. (**b**) Total number of protein data points for plexDIA, LF-DIA, and mTRAQ DDA at 1 % global protein FDR. (**c**) Venn-diagrams of each replicate for plexDIA and LF-DIA display protein groups quantified across samples A, B, and C. The mean number of proteins groups intersected across samples A, B, and C is 6,282 for plexDIA and 5,851 for LF-DIA. (**d**) The similarity between the quantified proteins across samples is quantified by the corresponding pairwise Jaccard indices to display data completeness. (**e**) Distributions of missing data for protein groups between pairs of runs of either the same sample (i.e., replicate injections) or between different samples. All analysis used match between runs. The corresponding results for the V2 duty cycle are shown in Fig. S3.

### plexDIA increases data completeness across samples

Next, we sought to compare LF-DIA and plexDIA in term of the consistency of protein quantification across samples. The systematic acquisition of ions by DIA is well established as a strategy for increasing the repeatability of peptide identification relative to shotgun DDA^24^. In addition to consistent data acquisition, plexDIA may further reduce the variability between samples and runs, and thus further increase the consistency (overlap) between quantified proteins relative to LF-DIA.

Indeed, both SILAC and isobaric labeling reduce missing data by enabling the quantification of peptides identified in at least one sample from a labeled set^18, 38^. Similarly, plexDIA takes advantage of the precisely known mass-shifts in the mass spectra for a peptide labeled with different tags to propagate peptide sequence identifications within a run. Specifically, confidently identified precursors in one channel (label) are matched to corresponding precursors in the other channels. This is the default analysis used with standards A, B and C. plexDIA has an additional mode for the special case when some proteins are present only in some samples of labeled sets. In such cases, plexDIA can enable sample specific identification for each protein by using multiple MS1- and MS2-based features to rigorously evaluate the spectral matches within a run and explicitly assign confidence for the presence of each protein in each sample. Such a special case is exampled by a plexDIA set in which one sample has both yeast and bacterial proteins while another sample has only yeast proteins, Fig. S2. These new analytical capabilities are described in the methods.

To assess whether plexDIA can improve data completeness, the protein groups intersecting across samples A, B and C were plotted as Venn diagrams for each replicate of plexDIA and LF-DIA, Fig. 2c. On average, the protein groups quantified in common across samples A, B, and C, were 6,282 for plexDIA and 5,851 for LF-DIA. The corresponding numbers for the V2 method are 7,923 for plexDIA and 8,318 for LF-DIA (Fig. S3c). Thus, a 3-plex plexDIA increased the rate of quantifying protein ratios across all 3 samples by 3.22 fold for the V1 method and by 2.86 fold for the V2 method, per unit time.

We further benchmarked the consistency of identified proteins both from the repeated analysis of the same sample (such as replicate injections of sample A) and from the analysis of different samples (such as comparing samples B and C). Consistent with prior reports for DIA data completeness, both LF-DIA and plexDIA identified largely the same proteins from replicate injections, quantified by high Jaccard indices and only about 13-15 % non-overlapping proteins, as shown in Fig. 2d,e. This overlap is comparable to the overlap of a high-quality LF-DIA dataset by Navarro, et al.^23^ as shown in Fig. S4. The overlap between the proteins identified in distinct samples remained similarly high for plexDIA while it was significantly reduced for the LF-DIA analysis, Fig. 2d,e. This increased repeatability for plexDIA likely arises from the fact that samples A, B, and C are analyzed in parallel as part of one set; this confers a further benefit of reduced missing data rate within a plexDIA set of only 2-3%, Fig. 2d,e. The larger the difference in protein composition between two samples, the higher the fraction of missing data for LF-DIA. In contrast, the missing data for plexDIA was low across all pairs of samples, Fig. 2e. The advantages of improved data completeness by plexDIA is especially pronounced when comparing the number of protein ratios from plexDIA and LF-DIA for samples which differ more in protein abundance, e.g. B and C; sample C has 6-fold more *E. coli* and 6-fold less *S. cerevisiae* relative to sample B. As a result, LF-DIA allowed to quantify only 1,383 ratios between *E. coli* and *S. cerevisiae* proteins while plexDIA allowed to quantify 1,807 protein ratios, Fig. 3a-c.

**Figure 3.**
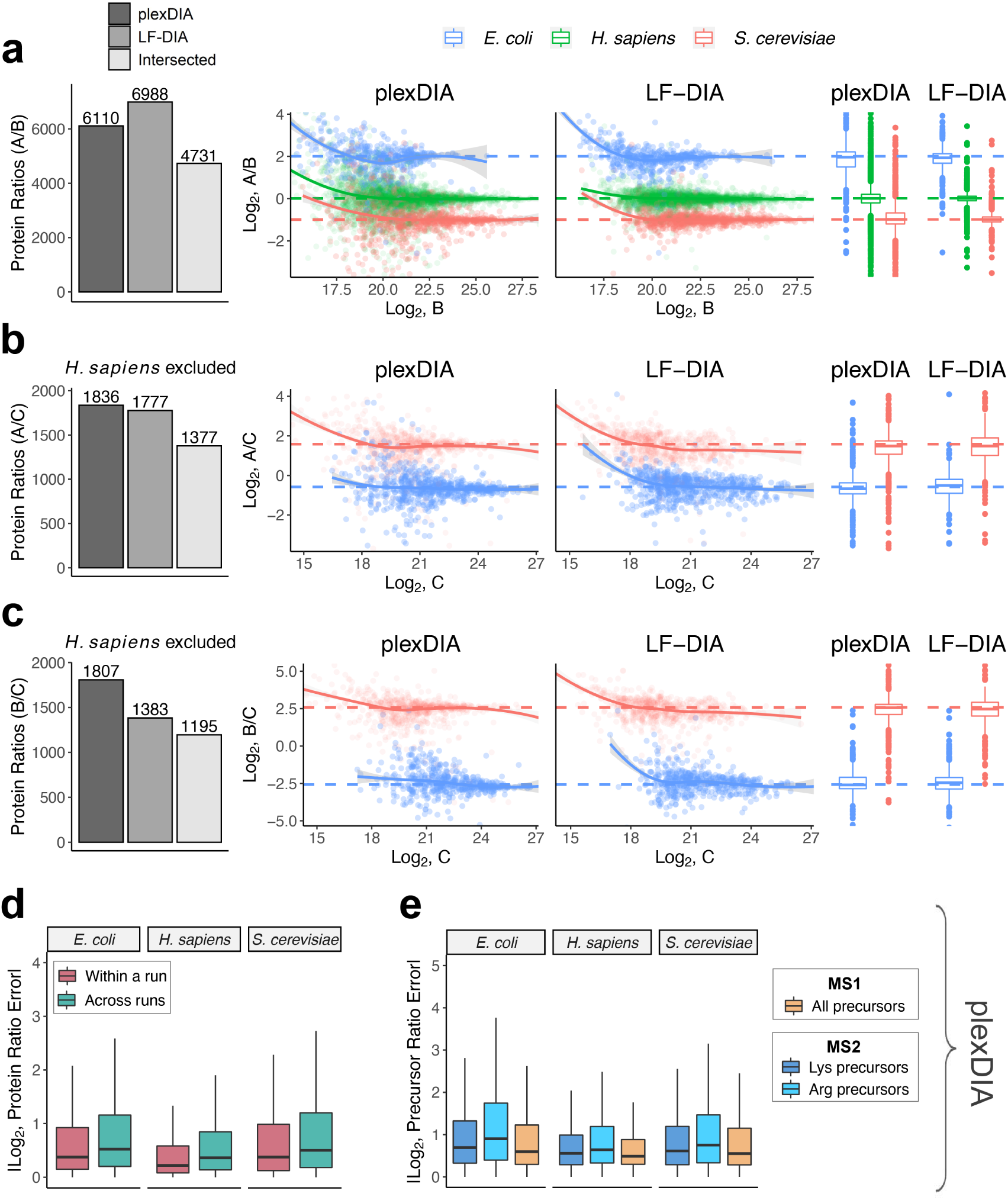
Quantitative accuracy and precision of plexDIA and LF-DIA. (**a**) Bars correspond to the number of quantified protein ratios between samples A and B by plexDIA, by LF-DIA, or by both methods (intersected proteins). To improve visibility, the scatter plot x and y axes were set to display data-points between 0.25% and 99.75% range. (**b**) Same as (**a**), but for samples A and C. (**c**) Same as (**a**), but for samples B and C. The protein ratios displayed in panels a-c are estimated from a single replicate, and two more replicates are shown in Fig. S6. (**d**) A comparison between the errors within and across plexDIA sets indicates similar accuracy. The error is defined as the difference between the mixing and the measured protein ratios for all pairs of samples, A/B, A/C, and B/C. The absolute values of these errors are displayed for samples within a plexDIA set (e.g., run2 A / run2 B) and for samples across sets (e.g., run1 A / run2 B). The corresponding accuracy within and across plexDIA for the V2 methods is shown in Fig. S5. (**e**) Absolute precursor ratio errors were calculated for samples A/B, A/C, and B/C and combined to compare ratio errors for MS1 and MS2 quantification. The MS2 quantification of precursors having C-terminal lysine or arginine is shown separately.^10^

### Quantitative accuracy of plexDIA is comparable to LF-DIA

To benchmark the quantitative accuracy and precision of plexDIA and LF-DIA, we compared the measured protein ratios between pairs of samples to the ones expected from the study design, Fig. 1. Because each sample contains a known amount of *E. coli*, *S. cerevisiae*, and *H. sapiens* protein lysate and most peptides are unique to each species, the protein ratios between pairs of samples correspond to the corresponding mixing ratios^23, 24^. The expected ratios allow for rigorous benchmarking of the accuracy and precision of plexDIA and LF-DIA. *H. sapiens* protein group ratios were excluded from analyses involving sample C as it would compare U-937 (A and B) to Jurkat (C) cell lines - therefore, deviations from expected ratios would be a combination of quantitative noise and cell-type specific differences in protein abundance.

For well controlled comparisons between the quantitative accuracy of LF-DIA and plexDIA, we used the set of protein ratios quantified by both methods. The comparison results from V1 are shown in Fig. 3a-c and from V2 in Fig. S5. These results indicate that on average plexDIA has comparable accuracy and precision to LF-DIA. Consistent with the expectation that labeling helps to control for nuisances, the results indicate that plexDIA quantification within a set is slightly more accurate than across sets, Fig. 3d. However, the difference is small, and accuracy across different plexDIA sets is high, Fig. S7a-c.

By design, plexDIA allows quantifying precursors based on MS2- and MS1-level data, and we evaluated the quantitative accuracy for both levels of quantification, Fig. 3e. Since both lysine and N-terminal amine groups are labeled by the amine-reactive mTRAQ labels, both b- and y-fragment ions of lysine peptides are sample-specific and thus contribute to MS2 level quantification. In contrast, only b-ions are sample-specific for arginine peptides, and thus only b-ions are used for their MS2-level quantification. As a result, the MS2-level quantification accuracy for arginine peptides is slightly lower, Fig. 3e. The small magnitude of this difference is likely attributable to the fact that mTRAQ stabilizes b-ions^39^. The accuracy of MS1-quantification by V1 is high for all peptides and slightly higher than the accuracy of MS2 quantification Fig. 3e. The MS2 optimized duty cycle (V2) resulted in deeper proteome coverage and lower accuracy for both LF-DIA and plexDIA, Fig. S5. However, different duty cycles implemented on different instruments will likely improve the accuracy and coverage by MS2-optimized methods.

### Repeatability of plexDIA is comparable to LF-DIA

To assess the repeatability of plexDIA and LF-DIA quantitation, we computed the coefficient of variation (CV) for proteins quantified in triplicate runs for each method using MaxLFQ abundances^40^; we required each protein group to be quantified three times for plexDIA and LF-DIA, then the CVs for the overlapping sample-specific protein groups (*n*=12,863) were plotted in Fig. 4a. The results indicate that plexDIA and LF-DIA have relatively consistent quantitation and comparable quantitative repeatability, with median CVs for repeated injections of 0.103 and 0.108, respectively. Repeatability of plexDIA was also compared for triplicates of the same labeled samples, and for triplicates in which each replicate had samples with alternating labels. Median CVs for the triplicates were 0.110 and 0.148 for ’same labels’ and ’different labels’ experiments, Fig. S7d.

**Figure 4.**
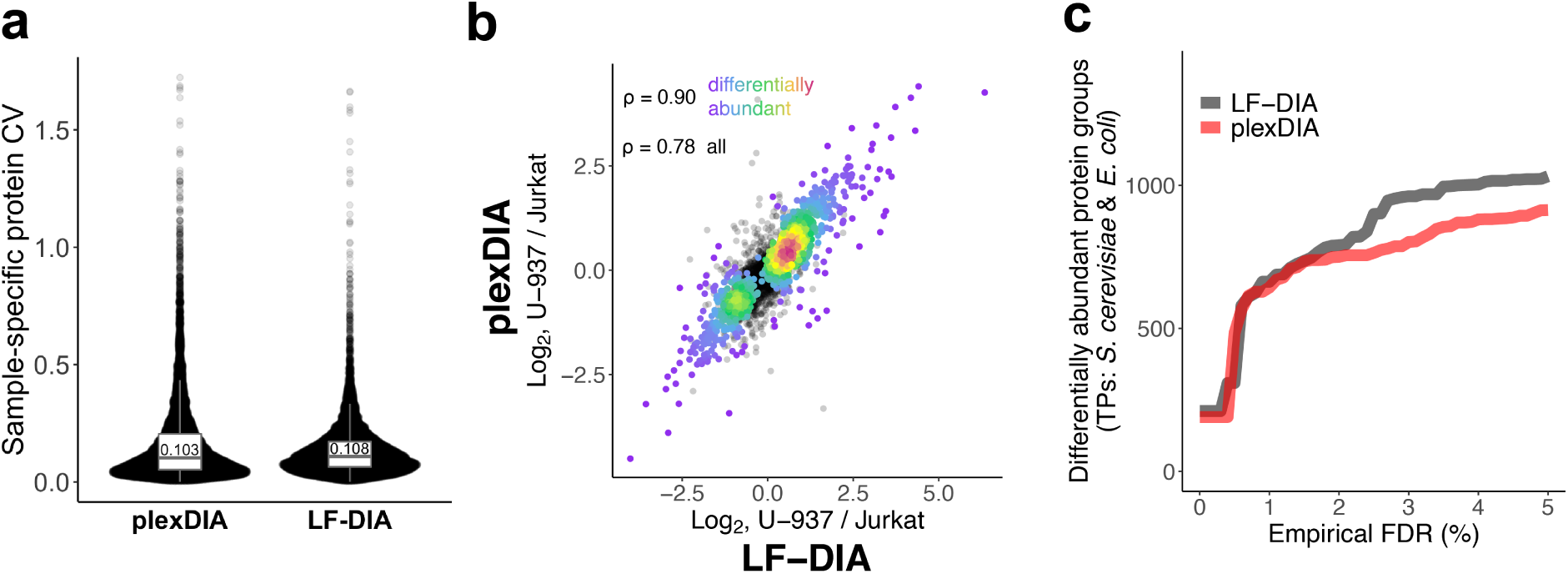
Using plexDIA to estimate differential protein abundance. (**a**) Quantitative repeatability was estimated by calculating coefficients of variation (CV) for MaxLFQ protein abundances (12,863 sample-specific protein data-points) calculated across triplicates for plexDIA and LF-DIA. (**b**) Proteins found to be differentially abundant between U-937 and Jurkat cells by LF-DIA were plotted as ratios of U-937/Jurkat for plexDIA and LF-DIA and colored by density. The Spearman correlation shown was calculated to quantify the agreement between the estimated relative protein levels of differentially abundant proteins at 1% FDR. Non-differentially abundant proteins are plotted in black; the Spearman correlation of all proteins (*n*=2,728) and differentially abundant proteins at 1% FDR (*n*=1,078) is 0.78 and 0.90, respectively. (**c**) Number of differentially abundant proteins between samples A and B as a function of the empirical FDR. The y-axis shows the number of true positives (only *S. cerevisiae* and *E. coli* proteins, which are differentially abundant) and the x-axis shows the false discovery rates estimated from the human proteins identified to be differentially abundant. The differential abundance was estimated using 3 replicates from each method, and thus LF-DIA took 3-times more instrument time per sample than plexDIA.

### Estimating differential protein abundance by plexDIA and LF-DIA

We investigated the agreement of differential protein abundance between U-937 and Jurkat cell lines with plexDIA and LF-DIA. Differential protein abundance was estimated from LF-DIA data, and the differentially abundant proteins at 1% FDR were used to assess the agreement between U-937 and Jurkat protein ratios estimated by plexDIA and LF-DIA, Fig. 4b. The estimates by the two methods are similar, as indicated by a Spearman correlation of 0.90 for differentially abundant proteins (*n*=1,078 at 1% FDR), and a Spearman correlation of 0.78 for all intersected human proteins (*n*=2,728) (Fig. 4b).

We also compared the ability of plexDIA and LF-DIA to recall true differentially abundant proteins as a function of each method’s empirical FDR. Our experimental design from Fig. 1c provides strong ground truth. It dictates that between samples A and B, only *S. cerevisiae* and *E. coli* are differentially abundant because they were spiked in at different ratios (1:2 and 4:1, respectively) while human proteins are not because they are present in a 1:1 ratio and compare the same cell type (U-937 monocytes). Therefore, true positives (*S. cerevisiae* and *E. coli* proteins) and true negatives (*H. sapiens* proteins) are known. With this prior knowledge, we compared the number of true positives for LF-DIA and plexDIA as a function of the empirical FDR, Fig. 4c. Both methods used 3 replicates and performed comparably at 1% empirical FDR, with 643 proteins and 663 proteins found to be differentially abundant for plexDIA and LF-DIA, respectively. The slight increase of true positives for LF-DIA at higher empirical FDR may be due to its slightly higher precision as visible in Fig. 3. In conclusion, plexDIA achieved comparable statistical power as LF-DIA while using 3-times less instrument time and expense.

### Cell division cycle analysis with plexDIA

Next, we applied plexDIA to quantify protein abundance across the cell division cycle (CDC) of U-937 monocyte cells. The CDC analysis allows further validation of plexDIA based on well established biological processes during the CDC while simultaneously offers the possibility of new discoveries. The ability of plexDIA to analyze small samples made it possible to isolate cells from different phases of the CDC based on their DNA content, Fig. 5a. The cell isolation used fluorescence activated cell sorting (FACS), which allowed us to analyze cell populations from G1, S, and G2/M phases without the artifacts associated with blocking the CDC to achieve population synchronization^41^.

**Figure 5.**
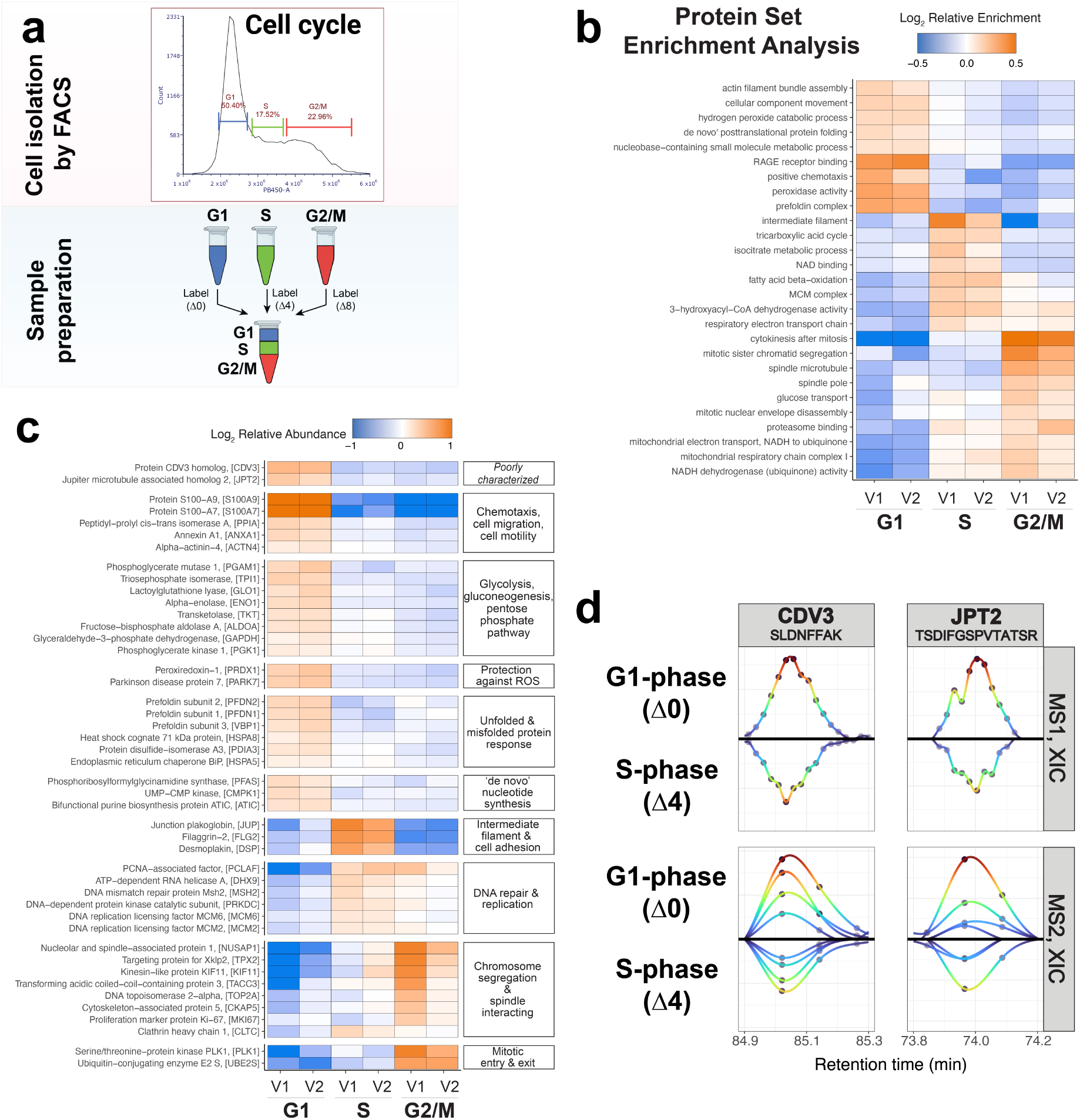
Cell cycle analysis with plexDIA. (**a**) U-937 monocytes were sorted by FACS based on DNA content to separate into G1, S, and G2/M cell-cycle phases; the samples were prepared as a plexDIA set, then analyzed with MS1 and MS2-optimized data acquisition methods (referred to as V1 and V2, respectively). (**b**) Protein set enrichment analysis of cell-cycle phases from plexDIA data. (**c**) A subset of proteins found to be differentially abundant at 1% FDR across cell-cycle phases were grouped by function, then plotted to show the relative abundances across cell-phases. (**d**) Extracted-ion chromatograms (XIC) at MS1 and MS2 for precursors from poorly characterized proteins, CDV3 and JPT2.

The peptides from the sorted cells were labeled with non-isobaric isotopologous labels, combined, and analyzed both by MS1-optimized (V1) and MS2-optimized (V2) plexDIA methods, Fig. 5a. By using different data acquisition methods, we aimed to (i) reduce systematic biases that may be shared by technical replicates and (ii) evaluate the agreement between MS1 and MS2-based quantification by plexDIA in the context of a biological experiment. Analyzing the V1 and V2 data with DIA-NN resulted in 4,391 unique protein groups and 4,107 gene groups at 1% global FDR. These data were filtered to include only proteotypic peptides, then gene-level information was used for downstream protein-set enrichment analysis (PSEA) and differential protein abundance analysis.

To identify biological processes regulated across the phases of the CDC, we performed PSEA using data from both V1 and V2, Fig. 5b. The V1 and V2 data indicated very similar PSEA patterns and identified canonical CDC processes, such as the activation of the MCM complex during S phase, and chromatid segregation and mitotic nuclear envelope disassembly during G2/M phase, Fig. 5b. These expected CDC dynamics and the agreement between V1 and V2 results demonstrate the utility of plexDIA for biological investigations. Furthermore, the PSEA indicated metabolic dynamics in the tricarboxylic acid (TCA) cycle and fatty acid beta-oxidation. These results provide direct evidence for the suggested coordination among metabolism and cell division^42, 43^.

To further explore the proteome remodeling during the CDC, we identified differentially abundant proteins across G1, S, and G2/M phase, Fig. 5c. From the 4,107 proteins identified across V1- and V2-acquired data, 400 proteins were found to be differentially abundant between cell cycle phases at 1% FDR. Some of these proteins are displayed in Fig. 5c organized thematically based on their functions. Consistent with results from PSEA, we find good agreement between V1 and V2 and expected changes in protein abundance, such as polo-like kinase 1 and ubiquitin-conjugating enzyme E2 peaking in abundance during G2/M phase.

In addition to the differential abundance of classic CDC regulators, we find that some poorly characterized proteins are also differentially abundant, such as proteins CDV3 and JPT2. To further investigate these proteins, we examined the extracted ion current (XIC) for representative peptides from these proteins, Fig. 5d. The XIC demonstrate consistent quantitative trends and coelution among precursors and peptide fragments labeled with different mass tags. This consistency among the raw data bolsters the confidence in new few findings by plexDIA, such as differential abundance of CDV3 and JPT2.

### Single-cell analysis with plexDIA

Next, we evaluated the potential of plexDIA to quantify proteins from single human cells. Thus, we prepared plexDIA sets from single cells from melanoma (WM989-A6-G3), pancreatic ductal adenocarcinoma (PDAC), and monocytes (U-937) cell lines were prepared into plexDIA sets using the nano-ProteOmic sample Preparation (nPOP)^44^.

We aimed to test its generalizability to different types of MS detectors, an orbitrap and a TOF detector, and its ability to take advantage of ion mobility technology, such as trapped ion mobility spectrometry^45^. The technologies were implemented by analyzing single-cell plexDIA samples using two commercial platforms, timsTOF SCP (Fig. 6a-f) and Q-Exactive classic (Fig. 6g- l). Both platforms achieved high quantitative accuracy and data completeness. To support high sample-throughput, both platforms used short chromatographic gradients to separate the peptides (Fig. 6d,j), which in the case of timsTOF SCP allowed quantifying about 1,000 proteins per cell while using about 10min of total instrument time (only 5min of active gradient) per single cell. Thus, plexDIA increases sample throughput by 3-12 fold over the top performing single-cell proteomics methods that do not utilize isobaric mass tags^46, 47^.

**Figure 6.**
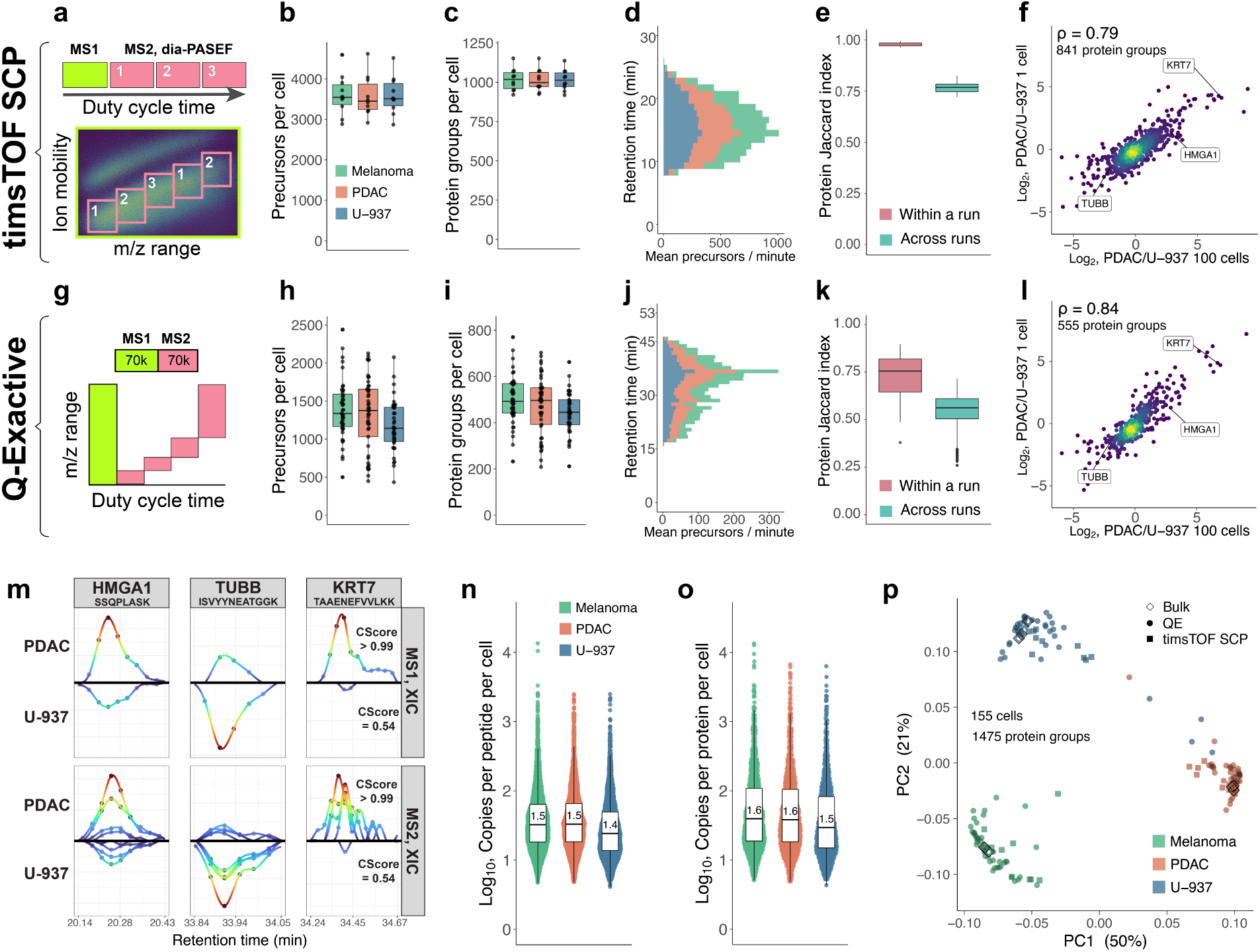
Single-cell protein analysis with plexDIA. (**a**) Cartoon visualizing the duty cycle used to analyze single-cell plexDIA sets with timsTOF SCP. Number of precursors (**b**) and protein groups (**c**) identified per single-cell. (**d**) Mean number of precursors identified from each cell-type per minute of chromatographic gradient. (**e**) Data completeness measured by Jaccard index within and between plexDIA sets. (**f**) Comparison of protein fold changes estimated from bulk samples (100 cells, x-axis) or single cells (y-axis). Panels (**g-l**) show analogous results to (**a-f**) but for data from a Q-Exactive classic. (**m**) Extracted-ion chromatograms (XIC) for precursors (MS1 level) and for peptide fragments (MS2 level) for peptides from differentially abundant proteins, HMGA1, TUBB, and KRT7; data is from single cells analyzed by Q-Exactive. Median number of copies for each peptide (**n**) and protein group (**o**), per single-cell; data is from single-cells analyzed by Q-Exactive. (**p**) Principal component analysis of 155 single-cells, including the cells analysed by timsTOF SCP or by Q-Exactive. The single cells are projected together with plexDIA triplicates of 100-cell bulk samples analyzed by Q-Exactive. All peptides and proteins shown are at 1% FDR.

As observed with bulk samples, plexDIA resulted in high data completeness among single-cell proteomes, Fig. 6e,k. It exceeded 98 % within labeled sets analyzed by timsTOF SCP (Fig. 6e) and remained over 50 % even between plexDIA sets analyzed by Q-Exactive, Fig. 6k. This high- level of data completeness is enabled by leveraging the co-elution of isotopologues with precisely known mass offsets, Fig. 1b. Still, about 5% of the single cells had comparable missing data to negative controls and were removed from downstream analysis as sample-preparation likely failed, Fig. S8.

plexDIA quantified protein fold-changes spanning a 1,000-fold dynamic range and exhibited good agreement with corresponding fold-changes quantified from 100-cell bulk samples, Fig. 6f,l.

To explore the raw data supporting these measurements, we plotted both MS1-level and MS2- level extracted ion current from pairs of isotopologous precursors, Fig. 6m. The data indicate that 1) the isotopologously labeled precursors co-elute and apex synchronously, and 2) the two lowly abundant precursors whose identification depended on the plexDIA module have precursors, fragments and intensities in excellent agreement with the more abundant isotopologues, and with the bulk measurements, Fig. 6l,m. These findings demonstrate that plexDIA may improve the sensitivity of single-cell proteomic analysis and thus increase data completeness, especially across cells with very different proteomes.

Sampling and detecting a sufficient number of precursor copies is key for accurate precursor quantification, otherwise quantification accuracy is undermined by counting noise^19, 48^. Since peptide fragmentation is usually incomplete, approaches like plexDIA that can perform MS1-level quantification are likely to count more copies per peptide than approaches relying on MS2 or MS3 level quantification^6^. To evaluate this expectation, we estimated the number of peptide and protein copies that the orbitrap counted from single cells, Fig. 6n,o. The estimates rely on orbitrap physics^49, 50^ and were not extended to the single-cell measurements by the timsTOF SCP.

Single-cell plexDIA data acquired from Q-Exactive and timsTOF SCP instruments were projected using a weighted PCA, Fig. 6p. To evaluate if the cell type separation is consistent with relative protein levels measured in bulk samples, we also projected 100-cell bulk plexDIA standards acquired on Q-Exactive, Fig. 6p. We found strong agreement between single-cell samples and 100-cell bulk samples. Similarly, single-cell data acquired by Q-Exactive and timsTOF SCP clustered by cell type, not platform type. To ensure that clustering was not an artifact of label-specific biases, we plotted the same PCA, except colored by the mTRAQ label that was used for tagging each single-cell and found little to no dependence of labels on clustering, Fig. S9.

## Discussion

While multiple methods allow increasing proteomics throughput, plexDIA is distinct in simultaneously allowing high sensitivity, depth and accuracy. plexDIA enables a multiplicative increase (3-fold with 3 labels) in the rate of consistent protein quantification across limited sample amounts while preserving proteomic coverage, quantitative accuracy, precision, and repeatability of LF-DIA. Similar to other labeling methods, such as TMT-DDA, parallel analysis of multiple samples by plexDIA saves LC-MS/MS time and costs. Currently, the commercially available labels for plexDIA are low-plex (mTRAQ, TMT0/TMT/shTMT, or dimethyl labeling^12^), compared to 18- plex isobaric TMTpro labels available for DDA^4^. This current plex disadvantage is offset by the parallel precursor analysis enabled by plexDIA. Indeed, quantifying about 8,000 proteins / sample took 0.5h for 3-plexDIA (Fig. S5) and 1.1h for a highly-optimized 16-plex TMTpro workflow^51^. Furthermore, 3-plexDIA affords higher sensitivity since it does not require offline fractionation and does not incur associated sample losses. We expect that the plexDIA framework will motivate the development of higher plex mass tags for plexDIA that are optimized for different applications, such as for single-cell proteomics^7^.

The parallel sample and peptide analysis by plexDIA becomes increasingly important for lowly abundant samples since they require long ion accumulation times that undermine the throughput of serial acquisition methods, such as TMT-DDA, even when the vast majority of MS2 scans result in confident peptide identifications^7, 52^. Thus, plexDIA is particularly attractive for the analysis of nanogram samples as it may afford accurate and deep proteome quantification without using 2-dimensional peptide separation (offline fractionation). Indeed, plexDIA is motivated in part by ideas for achieving sensitive and multiplexed single-cell proteomics^7, 19, 28^.

Our data demonstrate that plexDIA reduces the amount of missing data between diverse samples both within and across runs. This reduction stems from buffering sample-to-sample variability in protein composition. Furthermore, we introduced an approach for matching precursors within a run, which reduce missing data to mere 2-3 % in bulk samples (Fig. 2) and 2 % in single-cell samples (Fig. 6). Thus, plexDIA analysis of samples with variable protein composition or abundance results in less missing data. This opens the potential for further gains. For example, small samples could be labeled then combined with a labeled carrier sample to improve proteomic coverage of the smaller samples. Such nonisobaric carrier design will naturally extend the isobaric carrier concept^20, 28, 53^ and its benefits to DIA analysis to deep single-cell proteomics analysis. Indeed, the dynamic range, accuracy, and data completeness of the single-cell protein data obtained by plexDIA (Fig. 6) can enable interpreting natural variation across the proteomes of single cells^54^. plexDIA offers a framework that may scale to *n* labels, and thus increase throughput *n*-fold, reduce costs nearly *n*-fold, and increase the fraction of proteins quantified across all samples. Crucially, plexDIA maintains accurate quantification and good repeatability. Here, we demonstrate this potential for *n* = 3. Increasing *n* beyond 3 offers clear benefits but also faces challenges. One challenge is the increased potential for interference, which can be resolved by increasing the resolving power of MS scans and improving data analysis. Another challenge is sampling enough ions from each peptide given the finite capacity of MS detectors, which can be relieved by sampling smaller m/z ranges, e.g., quantification relying on small MS2 windows or split m/z ranges at MS1. The capacity of MS detectors is less limiting for small samples, such as single cells, and thus increasing the number of labels holds much potential for single-cell proteomics, as previously discussed^28, 55^.

The plexDIA framework intentionally uses nonisobaric isotopologous labels, which results in sample-specific precursors (allowing MS1 quantification) and in sample-specific peptide fragments (allowing MS2 quantification). Therefore, the plexDIA strategy enables quantification at the MS1 and MS2 levels, which offers advantages, such as evaluation of measurement reliability^56^. This strategy is opposite to previous approaches^33, 34^ and could in principle increase interference. Yet, this theoretical potential is effectively thwarted by our data analysis (Fig. 1b), and thus it does not significantly affect our results presented in Fig. 3.

In this work, we demonstrated the capabilities of plexDIA in providing a fold-change throughput increase for DIA proteomics, while yielding comparable data quality. While the 3-fold speed increase is already enabling for many applications, plexDIA unleashes opportunities beyond sample-throughput. For example, plexDIA can enable further gains in sensitivity for single-cell proteomics^7^, beyond the results demonstrated in Fig. 6. This may be achieved by including an isotopologous carrier channel, wherein a concentrated standard or pooled sample is used (i) to increase the sensitivity and thus identification numbers and data completeness in other channels, and (ii) to provide a reference signal for quantification. The quantitative aspect here has a double benefit. Quantification accuracy and robustness can be improved by (i) using MS1- and MS2-level signals that are minimally affected by interference and by (ii) calculating quantities relative to the internal standard, which is likely to also significantly reduce the batch effects associated with LCMS performance variation. This makes the technology introduced by plexDIA highly promising not just for very deep profiling of selected samples using offline fractionation, but also for large-scale experiments, wherein batch effects are a significant challenge. Another avenue of plexDIA is increasing the throughput of applications seeking to quantify protein interactions, conformations and activities. For example, plexDIA is readily compatible with the recently reported covalent protein painting that enables analysis of protein conformations in living cells^57, 58^. Since there are no fundamental limitations preventing the creation of non-isobaric labels which would allow a higher degree of multiplexing with DIA, we expect plexDIA to enable even higher throughput in the future. Given these considerations, we believe that plexDIA will eventually become the predominant DIA workflow, preferable over label-free approaches for most applications.

## Methods

### Cell culture

U-937 (monocytes) and Jurkat (T-cells) were cultured in RPMI-1640 Medium (Sigma-Aldrich, R8758), HPAF-II cells (pancreatic ductal adenocarcinoma (PDAC) cells, ATCC CRL-1997) were cultured in EMEM (ATCC 30-2003); all three cell-lines were supplemented with 10% fetal bovine serum (Gibco, 10439016) and 1% penicillin-streptomycin (Gibco, 15140122) and grown at 37*^◦^*C. Melanoma cells (WM989-A6-G3, a kind gift from Arjun Raj, University of Pennsylvania) were grown as adherent cultures in TU2% media which is composed of 80% MCDB 153 (Sigma-Aldrich M7403), 10% Leibovitz L-15 (ThermoFisher 11415064), 2 % fetal bovine serum, 0.5% penicillin-streptomycin and 1.68mM Calcium Chloride (Sigma-Aldrich 499609). All cells were harvested at a density of 10^6^ cells/mL and washed with sterile PBS. For bulk plexDIA benchmarks, U-937 and Jurkat cells were resuspended to a concentration of 5 *×* 10^6^ cells/mL in LC-MS water and stored at *−*80*^◦^*C.

*E. coli* and *S. cerevisiae* were grown at room-temperature (21*^◦^*C) shaking at 300 rpm in Luria Broth (LB) and yeast-peptone-dextrose (YPD) media, respectively. Cell density was measured by OD600 and cells were harvested mid-log phase, pelleted by centrifugation, and stored at *−*80*^◦^*C.

### Preparation of bulk plexDIA samples

The harvested U-937 and Jurkat cells were heated at 90*^◦^*C in a thermal cycler for 10 min to lyse by mPOP^59^. Tetraethylammonium bromide (TEAB) was added to a final concentration of 100 mM (pH 8.5) for buffering, then proteins were reduced in tris(2-carboxyethyl)phosphine (TCEP, Supelco, 646547) at 20 mM for 30 minutes at room temperature. Iodoacetamide (Thermo Scientific, A39271) was added to a final concentration of 15 mM and incubated at room temperature for 30 minutes in the dark. Next, Benzonase Nuclease (Millipore, E1014) was added to 0.3 units/*µ*L, Trypsin Gold (Promega, V5280) to 1:25 ratio of substrate:protease, and LysC (Promega, VA1170) to 1:40 ratio of substrate:protease, then incubated at 37*^◦^*C for 18 hours. *E. coli* and *S. cerevisiae* samples were prepared similarly; however, instead of lysis by mPOP, samples were lysed in 6 M Urea and vortexed with acid-washed glass beads alternating between 30 seconds vortexing and 30 seconds resting on ice, repeated for a total of 5 times.

After digestion, all samples were desalted by Sep-Pak (Waters, WAT054945). Peptide abundance of the eluted digests was estimated by nanodrop A280, and then the samples were dried by speed-vacuum and resuspended in 100 mM TEAB (pH 8.5). U-937, Jurkat, *E. coli*, and *S. cerevisiae* digests were mixed to generate three samples which we refer to as Sample A, B, and C, and the mixing ratios are described in Table S1. Samples A, B, and C were split into two groups: (i) was kept label-free, and (ii) was labeled with mTRAQ Δ0, Δ4, or Δ8 (SciEx, 4440015, 4427698, 4427700), respectively. An appropriate amount of each respective mTRAQ label was added to each Sample A-C, following manufacturers instructions. In short, mTRAQ was resuspended in isopropanol, then added to a concentration of 1 unit/100 *µ*g of sample and left to incubate at room-temperature for 2 hours. We added an extra step of quenching the labeling reaction with 0.25% hydroxylamine for 1 hour at room-temperature, as is commonly done in TMT experiments where the labeling chemistry is the same^6, 50^. After quenching, the mTRAQ-labeled samples (A-C) were pooled to produce the final multiplexed set used for benchmarking plexDIA.

### Preparation of single-cell plexDIA samples

Single cells were thawed from liquid nitrogen storage in 10 % DMSO and culture media at a concentration of 1 x 10^6^ cells/mL. Cells were first washed twice in PBS to remove DMSO and media and then were suspended in PBS at 200 cells/ *µ*L for sorting and sample preparation by nPOP as detailed by Leduc et al. [44]. In brief, single cells were isolated by CellenONE and prepared in droplets on the surface of a glass slide, including lysing, digesting, and labeling individual cells. In each droplet, single-cells were lysed in 100 % DMSO, proteins were digested with Trypsin Gold at a concentration of 120 ng/*µ*L and 5 mM HEPES pH 8.5, peptides were chemically labeled with mTRAQ, then finally single-cells were pooled into a plexDIA set for subsequent analysis. Cells were prepared in clusters of 3 for ease of downstream pooling into plexDIA sets; a total of 48 plexDIA sets were prepared per single glass slide.

Each plexDIA set was composed of a single PDAC, Melanoma, and U-937 cell, except if a negative control was present in place of a cell. For samples run on the Q-Exactive, every fourth set contained a negative control that received all the same reagents but did not include a single cell. This resulted in 132 single cells prepared with 12 total negative controls. 10 additional plexDIA sets were run on the timsTOF SCP for a total of 30 single cells (no negative controls). Celltypes were labeled with randomized mass tags designs in the plexDIA sets to avoid any systematic biases with labeling. Specifically, each cell type was labeled with each mass tag as described in the single-cell metadata file.

### Cell division cycle, FACS and sample preparation

U-937 monocytes were grown as described above, harvested and aliquoted to a final 1 mL suspension of approximately 1 *×* 10^6^ cells in RPMI-1640 Medium. Then DNA was stained by incubating the cells with Vybrant DyeCycle Violet Stain (Invitrogen, V35003) at a final concentration of 5 *µ*M in the dark for 30 minutes at 37*^◦^*C, as per the manufacturer’s instructions. Next, the cells were centrifuged, then resuspended in PBS to a density of 1 *×* 10^6^ cells/mL. The cell suspension was stored on ice and protected from light until sorting began.

The cells were then sorted with a Beckman CytoFLEX SRT. The population of U-937s was gated to select singlets based on FSC-A and FSC-H, this population of singlets was then subgated based on DNA content using the PB-450 laser (ex = 405 nm / em = 450 nm). The G1 population is the most abundant population in actively dividing cells, and the G2/M populations should theoretically have double the intensity (DNA content), while the S-phase lies in between. Populations of G1, S, and G2/M cells were collected based on these subgates and sorted into 2 mL Eppendorf tubes.

Post-sorting, the cells were centrifuged at 300g for 10 minutes, PBS was removed, then the cells were resuspended in 20 *µ*L HPLC water to reach a density of approximately 4,000 cells/*µ*L. The cell suspensions were lysed using the Minimal ProteOmic sample Preparation (mPOP) method, which involves freezing at -80*^◦^*C and then heating to 90*^◦^*C for 10 minutes^59^. Next, the cell lysates were prepared exactly as described in the “Sample Preparation” section. In brief, the cell lysate was buffered with 100 mM TEAB (pH 8.5), then proteins were reduced with 20 mM TCEP for 30 minutes at room temperature. Next, iodoacetamide was added to a final concentration of 15 mM and incubated at room temperature for 30 minutes in the dark, then Benzonase Nuclease was added to 0.3 units/*µ*L. Trypsin Gold and LysC to were then added to the cell lysate at 1:25 and 1:40 ratio of protease:substrate, then the samples were incubated at 37*^◦^*C for 18 hours. After digestion, the peptides were desalted by stage-tipping with C18 extraction disks (Empore, 66883-U) to remove any remaining salt that was introduced during sorting^60^. G1 cells were labeled with mTRAQ Δ0, S cells were labeled with mTRAQ Δ4, and G2/M cells were labeled with Δ8, then combined to form a plexDIA set of roughly 2,000 cells per cell-cycle phase (label). The combined set was analyzed with 2 hour active gradients of MS1 (V1) and MS2-optimized (V2) methods as described in the “Acquisition of bulk data” section.

### Acquisition of bulk data

Multiplexed and label-free samples were injected at 1*µ*L volumes via Dionex UltiMate 3000 UH- PLC to enable online nLC with a 25 cm × 75 *µm* IonOpticks Aurora Series UHPLC column (AUR2-25075C18A). These samples were subjected to electrospray ionization (ESI) and sprayed into a Thermo Q-Exactive orbitrap for MS analysis. Buffer A is made of 0.1% formic acid (Pierce, 85178) in LC-MS-grade water; Buffer B is made of 80 % acetonitrile and 0.1 % formic acid mixed with LC-MS-grade water.

The gradient used for LF-DIA is as follows: 4% Buffer B (minutes 0-11.5), 4%-5% Buffer B (minutes 11.5-12), 5%-28% Buffer B (minutes 12-75), 28%-95% Buffer B (minutes 75-77), 95% Buffer B (minutes 77-80), 95%-4% Buffer B (minutes 80-80.1), then hold at 4% Buffer B until minute 95, flowing at 200 nl/min throughout the gradient. The V1 duty cycle was comprised of 5x(1 MS1 full scan x 5 MS2 windows) as illustrated in Fig. 1b. Thus, the duty cycle has a total of 25 MS2 windows to span to full m/z scan range (380-1370 m/z) with 0.5 Th overlap between adjacent windows. The length of the windows was variable for each subcycle (20 Th for subcycles 1-3, 40 Th for subcycle 4, and 100 Th for subcycle 5). Each MS1 full scan was conducted at 140k resolving power, 3×10^6^ AGC maximum, and 500 ms maximum injection time. Each MS2 scan was conducted at 35k resolving power, 3×10^6^ AGC maximum, 110 ms maximum injection time, and 27% normalized collision energy (NCE) with a default charge of 2. The RF S-lens was set to 80%. The V2 duty cycle consisted of one MS1 scan conducted at 70k resolving power with a 300 ms maximum injection time and 3×10^6^ AGC maximum, followed by 40 MS2 scans at 35k resolving power with 110 ms maximum injection time and 3×10^6^ AGC maximum. The window length for the first 25 MS2 scans was set to 12.5 Th; the next 7 windows were 25 Th, then the last 8 windows were 62.5 Th. Adjacent windows shared a 0.5 Th overlap. All other settings were the same as the LF-DIA V1 method. All label-free samples for bulk benchmarking containing *S. cerevisiae*, *E. coli*, and *H. sapiens* were run in triplicate. However, the third run of LF-DIA sample C using the V2 method was an outlier and omitted from analysis due to poor performance.

mTRAQ labeling increases hydrophobicity of peptides, which is why a higher % Buffer B is used during the active gradient of multiplexed samples; in addition, the scan range was shifted 100 m/z higher than LF-DIA to account for the added mass of the label. The gradient used for plexDIA is as follows: 4% Buffer B (minutes 0-11.5), 4%-7% Buffer B (minutes 11.5-12), 7%-32% Buffer B (minutes 12-75), 32%-95% Buffer B (minutes 75-77), 95% Buffer B (minutes 77-80), 95%- 4% Buffer B (minutes 80-80.1), then hold at 4% Buffer B until minute 95, flowing at 200 nl/min throughout the gradient. The plexDIA V1 duty cycle was comprised of 5x(1 MS1 full scan x 5 MS2 windows), for a total of 25 MS2 windows to span to full m/z scan range (480-1470 m/z) with 0.5 Th overlap between adjacent windows. The length of the windows was variable for each subcycle (20 Th for subcycles 1-3, 40 Th for subcycle 4, and 100 Th for subcycle 5). Each MS1 full scan was conducted at 140k resolving power, 3×10^6^ AGC maximum, and 500 ms maximum injection time.

Each MS2 scan was conducted at 35k resolving power, 3×10^6^ AGC maximum, 110 ms maximum injection time, and 27% normalized collision energy (NCE) with a default charge of 2. The RF S-lens was set to 30%. The plexDIA V2 duty cycle consisted of one MS1 scan conducted at 70k resolving power with a 300 ms maximum injection time and 3×10^6^ AGC maximum, followed by 40 MS2 scans at 35k resolving power with 110 ms maximum injection time and 3×10^6^ AGC maximum. The window length for the first 25 MS2 scans was set to 12.5 Th; the next 7 windows were 25 Th, then the last 8 windows were 62.5 Th. Adjacent windows shared a 0.5 Th overlap. All other settings were the same as the plexDIA V1 method. Data acquired for the cell-division-cycle used 2 hour active gradients of the V1 and V2 methods.

The gradient used for mTRAQ DDA is the same used for plexDIA. However, the duty cycle was a shotgun DDA method. The MS1 full scan range was 450-1600 m/z, and was performed with 70k resolving power, 3×10^6^ AGC maximum, and 100 ms injection time. This shotgun DDA approach selected the top 15 precursors to send for MS2 analysis at 35k resolving power, 1×10^5^ AGC maximum, 110 ms injection time, 0.3 Th isolation window offset, 0.7 Th isolation window length, 8×10^3^ minimum AGC target, and 30 second dynamic exclusion.

### Acquisition of single-cell data

#### Q-Exactive

plexDIA single cell sets and 100-cell standards were injected at 1*µ*L volumes via Dionex UltiMate 3000 UHPLC to enable online nLC with a 15 cm × 75 *µm* IonOpticks Aurora Series UHPLC column (AUR2-15075C18A). These samples were subjected to electrospray ionization (ESI) and sprayed into a Thermo Q-Exactive orbitrap for MS analysis. Buffer A is made of 0.1% formic acid (Pierce, 85178) in LC-MS-grade water; Buffer B is made of 80 % acetonitrile and 0.1 % formic acid mixed with LC-MS-grade water. The gradient used is as follows: 4% Buffer B (minutes 0-2.5), 4%-8% Buffer B (minutes 2.5-3), 8%-32% Buffer B (minutes 3-33), 32%-95% Buffer B (minutes 33-34), 95% Buffer B (minutes 34-35), 95%-4% Buffer B (minutes 35-35.1), then hold at 4% Buffer B until minute 53, flowing at 200 nl/min throughout the gradient. The plexDIA duty cycle was comprised of 1 MS1 followed by 4 DIA MS2 windows of variable m/z length (specifically 120 Th, 120 Th, 200 Th, and 580 Th) spanning 378-1402 m/z. Each MS1 and MS2 scan was conducted at 70k resolving power, 3×10^6^ AGC maximum, and 300 ms maximum injection time. Normalized collision energy (NCE) was set to 27% with a default charge of 2. The RF S-lens was set to 80%.

To generate a spectral library from 100-cell standards on the Q-Exactive, the same settings were used with the exception that the duty consisted of 1 MS1 and 25 MS2 windows of variable m/z length (specifically 18 windows of 20 Th, 2 windows of 40 Th, 3 windows of 80 Th, and 2 windows of 160 Th). The MS2 scans were conducted at 35k resolving power, 3×10^6^ AGC maximum, and 110 ms maximum injection time.

#### timsTOF SCP

The single-cell plexDIA sets were separated on a nanoElute liquid chromatography system (Bruker Daltonics, Bremen, Germany) using a 25 cm × 75 *µm*, 1.6 *µm* C18 (AUR2- 25075C18A-CSI, IonOpticks, Au). The analytical column was kept at 50 °C. Solvent A was 0.1% formic acid in water, and solvent B was 0.1% formic acid in acetonitrile. The column was equilibrated with 4 column volumes of mobile phase A prior to sample loading. The peptides were separated over 30 min at 250 nL/min using the following gradients: from 2% to 17% B in 15 min, from 17% to 25% B in 5 min, 25% to 37% B in 3 min, 37%-85% B in 3 min, maintained at 85% for 4 min.

The timsTOF SCP was operated in dia-PASEF mode with the following settings: Mass Range 100 to 1700 m/z, 1/k0 Start 0.6 V s/cm2, End 1.2 V s/cm2, ramp and accumulation times were set to 166ms, Capillary Voltage was 1600V, dry gas 3 l/min, and dry temp 200 *^o^C*. dia-PASEF settings: Each cycle consisted of 1x MS1 full scan and 5x MS2 windows covering 297.7 – 797.7 m/z and 0.63 - 1.10 1/k0. Each window was 100 Th wide by 0.2 V s/cm2 high. There was no overlap in either m/z or 1/k0 (Fig. 6). The cycle time was 0.68 seconds. CID collision energy was 20 to 59eV as a function of the inverse mobility of the precursor.

### Spectral library generation

The *in silico* predicted spectral library used in LF-DIA analysis was generated by DIA-NN’s (version 1.8.1 beta 16) deep learning-based spectra and retention time (RT), and IMs prediction based on Swiss-Prot *H. sapiens*, *E. coli*, and *S. cerevisiae* FASTAs (canonical & isoform) downloaded in February 2022. The spectral library used for plexDIA benchmarking was created in a similar process, with the exception of a few additional commands entered into the DIA-NN command line GUI: 1) *{*–fixed-mod mTRAQ 140.0949630177, nK*}*, 2) *{*–original-mods*}*. Two additional libraries were generated: 1) mTRAQ-labeled spectral library from FASTAs containing only *E. coli*, and *S. cerevisiae* sequences. 2) mTRAQ-labeled spectral library from a FASTA containing only *H. sapiens* sequences; the former was used to search data shown in Fig. S2, and the latter was used to search cell-division-cycle and 100-cell standards. Triplicates of 100-cell standards of PDAC, Melanoma, and U-937 cells were run with the 1 MS1 x 25 MS2 scans method, searched using the *in silico*-generated human-only spectral library. The results of this search generated a sample-specific library covering about 5,000 protein groups; this library was used the search single-cell plexDIA sets acquired on the Q-Exactive and on the timsTOF SCP, as well as 100-cell standards run on the Q-Exactive with the same method used to acquire single-cell plexDIA data.

### plexDIA module in DIA-NN

A distinct feature of DIA-MS proteomics is the complexity of produced spectra, which are a mixture of fragments ions originating from multiple co-isolated precursors. This complexity has necessitated the rise of a variety of highly sophisticated algorithms for DIA data processing. Current DIA software, such as DIA-NN^25^, aims to find peak groups in the data that best match the theoretical information about such peptide properties as the MS/MS spectrum, the retention time and the ion mobility. Once identified correctly, the peak group, that is the set of extracted ion chromatograms of the precursor and its fragments in the vicinity of the elution apex, allows to integrate either the MS1- or MS2-level signals to quantify the precursor, which is the ultimate purpose of the workflow.

Similar to match-between-runs (MBR) algorithms, plexDIA data provide the opportunity to match corresponding ions, in this case between the same peptide labeled with different mass tags. However, the use of isotopologous mass tags, such as mTRAQ, allows to match the retention times within a run with much higher accuracy than what can be achieved across runs. Thus, the sequence propagation can be more sensitive and reliable than with MBR^7^. This allows to enhance sequence identifications analogously to the isobaric carrier concept introduced by TMT-based single-cell workflows^53, 61^. With the isobaric carrier approach, a carrier channel is loaded with a relatively high amount of peptides originating from a pooled sample that facilities peptide sequence identification^20, 28^. We implemented a similar approach in the plexDIA module integrated in DIA-NN. Once a peptide is identified in one of the channels, this allows to determine its exact retention time apex, which in turn helps identify and quantify the peptide in all of the channels by integrating the respective precursor (MS1) or fragment ion (MS2) signals.

Apart from the identification performance, plexDIA also can increase quantification accuracy.

The rich complex data produced by DIA promotes more accurate quantification because of algorithms that select signals from MS/MS fragment ions which are affected by interferences to the least extent^25^. For LF-DIA, DIA-NN selects fragments in a cross-run manner: fragments which tend to correlate well with other fragments across runs are retained, while those which often exhibit poor correlations due to interferences are excluded from quantification. While this approach yields good results, a limitation remains for LF-DIA: fragment ions only affected by interferences in a modest proportion of runs are still used for quantification, thus undermining the reliability of the resulting quantities in those runs. Here plexDIA provides a unique advantage. Theoretically, a single MS1- or MS2-level signal with minimal interference is sufficient to calculate the quantitative ratio between the channels. In this case, if low interference quantification is possible in at least one ‘best’ channel, this quantity can be multiplied by the respective ratios across other channels to obtain accurate estimates of quantities in all channels that share at least one low interference signal with this ‘best’ channel. This idea is implemented in DIA-NN to produce ‘translated’ quantities, which have been corrected by using ratios of high quality MS1 or MS2 signals between channels as described in Fig. 1b and Fig. S1.

### Data analysis with DIA-NN

DIA-NN (version 1.8.1 beta 16) was used to search LF-DIA and plexDIA raw files, which is available at plexDIA.slavovlab.net and scp.slavovlab.net/plexDIA. All LF-DIA benchmarking raw files were searched together with match between runs (MBR) if the same duty cycle was used; likewise, all plexDIA benchmarking raw files were searched together with MBR if the same duty cycle was used with the exception of the cell-division-cycle experiments which used V1 and V2 methods - these two runs were searched together.

DIA-NN search settings: Library Generation was set to “IDs, RT, & IM Profiling”, Quantification Strategy was set to “Peak height”, scan window = 1, Mass accuracy = 10 ppm, and MS1 accuracy = 5 ppm, “Remove likely interferences”, “Use isotopologues”, and “MBR” were enabled. Additional commands entered into the DIA-NN command line GUI for plexDIA: 1) *{*–fixed-mod mTRAQ 140.0949630177, nK*}*, 2) *{*–channels mTRAQ, 0, nK, 0:0; mTRAQ, 4, nK, 4.0070994:4.0070994; mTRAQ, 8, nK, 8.0141988132:8.0141988132*}*, 3) *{*–original-mods*}*, 4) *{*–peak-translation*}*, 5) *{*–ms1-isotope-quant*}*, 6) *{*–report-lib-info*}*, and 7) *{*–mass-acc-quant 5.0*}*. Note, #7 is only necessary for instances when MS2 quantitation is intended to be used; this command will use the pre-defined mass accuracy (e.g. 10 ppm) to identify precursors, but restrict the mass error tolerance to the value specified for quantitation; this can help reduce the impact of interferences for MS2-level quantitation. For LF-DIA, only the following additional commands were used: 1) *{*–original-mods*}*, 2) *{*–peak-translation*}*, 3) *{*–ms1-isotope-quant*}*, 4) *{*–report-lib-info*}*, and 5) *{*–mass-acc-quant 5.0*}*. The same search settings were used for single-cell Q-Exactive and timsTOF SCP data, however ’scan window’ was increased to 5.

### Data analysis with MaxQuant, DDA

MaxQuant (version 1.6.17.0) was used to search triplicate mTRAQ DDA, bulk benchmarking runs. MBR was enabled, and ‘Type’ was selected as ‘Standard’ with ‘Multiplicity’ = 3; mTRAQ-Lys0 & mTRAQ-Nter0, mTRAQ-Lys4 & mTRAQ-Nter4, and mTRAQ-Lys8 & mTRAQ-Nter8 were selected for light, medium, and heavy labels. Variable modifications included Oxidation (M), Acetyl (Protein-N-term); Carbamidomethyl (C) was selected as a fixed modification. Trypsin was selected as the protease, and searched with max. missed cleavage = 2.

### Quantifying proteins for bulk plexDIA benchmarks

MaxLFQ abundance for protein groups was calculated based on MS1 intensities (specifically the “MS1 Area” column output by DIA-NN) using the DIA-NN R package^25^ for data acquired with the V1 method. However, for data acquired using the V2 method, MS2 quantitation (specifically the “Precursor Translated” column output by DIA-NN) was used for quantitation. These protein abundances were used to calculate protein ratios across samples, which were normalized by sub- setting human proteins (which are present in a 1:1 ratio, theoretically) and multiplying by a scalar such that the human protein ratios were centered on 1, and thus the other species (*E. coli*, *S. cerevisiae*) would be systematically shifted to account for any small loading differences across samples.

The quantitative comparisons between LF-DIA and plexDIA throughout this article are for intersected sets of proteins so that the results would not be influenced by proteins analyzed only by one method and not the other. For examples, compared distributions were for the same set of proteins to avoid “survival biases”^62^.

### Protein-set enrichment analysis (PSEA)

PSEA was performed across the multiplexed bulk samples corresponding to cells sorted by DNA content into cell cycle phases (G1, S, and G2/M). The reference human gene set database was acquired from GOA^63^. The Kruskall Wallis test was used to determine whether the hypothesis that all multiplexed samples had equivalent median protein abundances for a functionally annotated group of proteins could be rejected at a q value *≤* 0.05. Only protein sets with at least 4 proteins present were statistically tested. PSEA was run separately for the multiplexed samples analyzed by V1 and V2 methods. Protein sets were combined from both data-acquisiton methods if at least one method produced a q value *≤* 0.05.

### Differential protein abundance testing

Differential protein abundance testing was performed using precursor-level quantitation. To account for variation in sample loading amounts, precursors from each sample were normalized to their sample-median. Then, each precursor was normalized by its mean across samples to convert it to relative levels. The normalized relative precursor intensities from different replicates were grouped by their corresponding protein groups and compared by a two-tailed t-test (Fig. 4b,c) or ANOVA (Fig. 5c) to estimate the significance of differential protein abundance across samples/conditions. This comparison captures both the variability between different replicates and different peptides originating from the same protein. To correct for multiple hypotheses testing, we used the Benjamini-Hochberg (BH) method to estimate q-values for differential abundance of proteins and protein sets.

### Relative protein fold-change between U-937 cells and Jurkat cells, bulk

Protein group abundances for were calculated by MaxLFQ from triplicates of LF-DIA and plex- DIA; specifically, sample B and sample C were compared to calculate relative fold-changes between *H. sapiens* cell-lines, U-937 and Jurkat. The protein groups plotted were required to be quantified in each of the triplicates of plexDIA and LF-DIA. A Spearman correlation was calculated for all protein groups and for differentially abundant protein groups.

### Correcting isotopic envelope of plexDIA precursors

mTRAQ labels, which were used in this demonstration of plexDIA, are separated by 4 Daltons (Da). Because C-terminal arginine precursors are singly-labeled and have a mere 4 Da separating isotopologous precursors, there is greater potential of isotopic envelope interference from lighter channels into heavier channels than there is for C-terminal lysine precursors which would be separated by 8 Da; therefore, to improve quantitative accuracy, we correct the theoretical super- position of isotopic envelopes between channels for C-terminal arginine precursors. This can be accomplished because each precursor has a well-defined theoretical distribution of isotopes that we model with a binomial distribution; we use this theoretical distribution of isotopes to subtract and add back a precise amount of signal from heavier channels to lighter channels for MS1-level quantitation of each precursor.

### Extracted ion current (XIC)

A precursor from a subset of proteins found to be differentially abundant was selected to be plotted to display the extracted ion current at MS1 and for fragments at MS2. Ion current was extracted using the DIA-NN GUI command interface by typing *{*–vis 25, PEPTIDE*}* where “PEPTIDE” is the peptide sequence and “25” is the number of scans to extract. MS1 and MS2 XICs were plotted to show the full elution profile. The four highest correlated fragments at MS2 were plotted; y-ions from C-terminal arginine peptide were excluded from plotting at MS2-level because these fragments are a super-position across samples as the C-terminus of arginine peptides is not labeled, and therefore, not sample-specific. The lines in Fig. 5d and Fig. 6m were colored dynamically as a function of intensity.

### Estimating peptide and protein copy numbers

Precursor copy numbers at the MS1-level were estimated based on the signal-to-noise level (S/N) of individual peaks. The noise level of centroided spectra were used as reported by the Thermo firmware and extracted using a modified version of the ThermoRawFileParser^64^. Precursors reported by DIA-NN were matched to the S/N data based on the reported retention time with a tolerance of 5 scans and 12 ppm mass error. The number of charges in an orbitrap is proportional to the S/N level and scales with a linear factor *C_N_* . This factor has been estimated to be *C_N_* = 3.5 for the Q-Exactive orbitrap^65, 66^ and has been confirmed by investigations with high-field orbitraps^49^. This proportionality constant was estimated at a resolving power of 240, 000 and must be scaled by the square root ratio with the resolving power used for acquiring the spectra (*R* = 70, 000). Precursor copy numbers are then calculated based on the number of charges *z* per precursor.

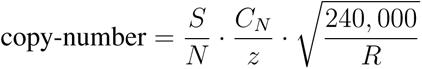

Analogous to the quantification, copy numbers were summed over the M and M+1 peaks. Peptide-level copy numbers were calculated as the sum of all charge states found for a given peptide; protein-level copy numbers were calculated as the sum of all peptides not shared with other proteins (proteotypic).

### Single cell data analysis

To increase sensitivity of single-cell analysis, Ms1.Extracted quantities output by DIA-NN were used for quantitation rather than Ms1.Area. Single cells with more than 60% missing data (no extracted MS1-level quantitation) at precursor-level were considered to have failed in sample preparation and were removed from analysis. Quantitative accuracy of single-cell sets was assessed by calculating fold-change between PDAC and U-937 cell-types of averaged single-cell MaxLFQ protein quantities and calculating a Spearman correlation to 100-cell bulk comparisons. The 100- cell bulk comparisons consisted of triplicates in which the each replicate alternated the labeling scheme. For a protein group to be included in the comparison, it was required to be quantified in at least 5 single-cells, and 2/3 of the bulk triplicates. Both the timsTOF SCP single-cell data and QE single-cell data were benchmarked to the same 100-cell QE-acquired plexDIA sets. Because missing data in DIA is related to low protein abundance, the missing MaxLFQ protein abundances in single cells and bulk were imputed with the lowest non-zero protein abundance for that protein in the same cell-type and condition (bulk or single-cells). The mean of each protein across the single cell observations and bulk triplicates (respectively) was taken to represent that cell-type and condition-specific protein abundance.

Single-cell sets acquired on the timsTOF SCP and QE were prepared on different days with different batches of cells. Generally, the data is quite similar as indicated by PCA Fig. 6p, but quantitative discrepancies between bulk samples which were acquired on the QE from one batch of cells, and single cell sets on the timsTOF SCP from another batch of cells may arise from real cellular differences as they were prepared from different cellular batches.

100-cell bulk plexDIA triplicates were used to identify proteins which are differentially abundant between U-937 and PDAC cells. Three proteins were chosen, and one precursor from each protein was selected to have its ion-current extracted and plotted from single-cell Q-Exactive acquired data. Please see the ”Extracted ion current (XIC)” subsection for more details about how this is performed.

PCA was performed on Ms1.Extracted timsTOF SCP single-cell, Q-Exactive single-cell, and Q-Exactive 100-cell data. The following is a brief outline of the computational workflow: the abundance of each precursor was divided by the mean abundance of all 3 isotopologous precursors within the plexDIA set; then, the precursors of each labeled cell in each plexDIA was normalized to its median abundance; then, each normalized precursor was divided by the mean of normalized precursor abundance across all labels and sets. These normalized precursor abundances were collapsed to protein group level by the median normalized abundance precursor. The protein group data was then normalized in the same way the precursors were normalized. Missing protein group data for each cell was imputed by K-nearest-neighbors; the data set was batch-corrected; and finally, a weighted PCA was generated from the data, as was previously described^50^.

### Data availability

The raw data and search results are available at MassIVE: MSV000089093

### Code availability

Data, code & protocols are available at plexDIA.slavovlab.net and github.com/SlavovLab/plexDIA. Supporting information for the single-cell plexDIA is available at: scp.slavovlab.net/plexDIA and scp.slavovlab.net/Derks et al 2022

## Acknowledgments

We thank A. Makarov, E. Gordon, and D. Perlman for discussions and constructive comments. This work was funded by a New Innovator Award from the NIGMS from the National Institutes of Health to N.S. under Award Number DP2GM123497, an Allen Distinguished Investigator award through The Paul G. Allen Frontiers Group to N.S., a Seed Networks Award from CZI CZF2019-002424 to N.S. This work received further support from the Francis Crick Institute, which receives its core funding from Cancer Research UK (FC001134), the UK Medical Research Council (FC001134), and the Wellcome Trust (FC001134 and IA 200829/Z/16/Z), as well as the European Research Council (SyG 951475 to M.R.). The work was further supported by the German Federal Ministry of Education and Research (BMBF), as part of the National Research Node ’Mass spectrometry in Systems Medicine’ (MSCoresys), under grant agreements 031L0220A to M.R. and 161L0221 to V.D..

## Author contributions

**Experimental design**: J.D., N.S. and V.D.

**LC-MS/MS**: J.D., M.W., S.K., G.H. and H.S.

**Sample preparation**: J.D. and A.L.

**Raising funding**: N.S., M.R. and V.D.

**Supervision**: N.S.

**Data analysis**: J.D., G.W., V.D., and N.S.

**Initial draft**: J.D., V.D., and N.S.

**Writing**: All authors approved the final manuscript.

## Supplementary Figures

**Figure S1.**
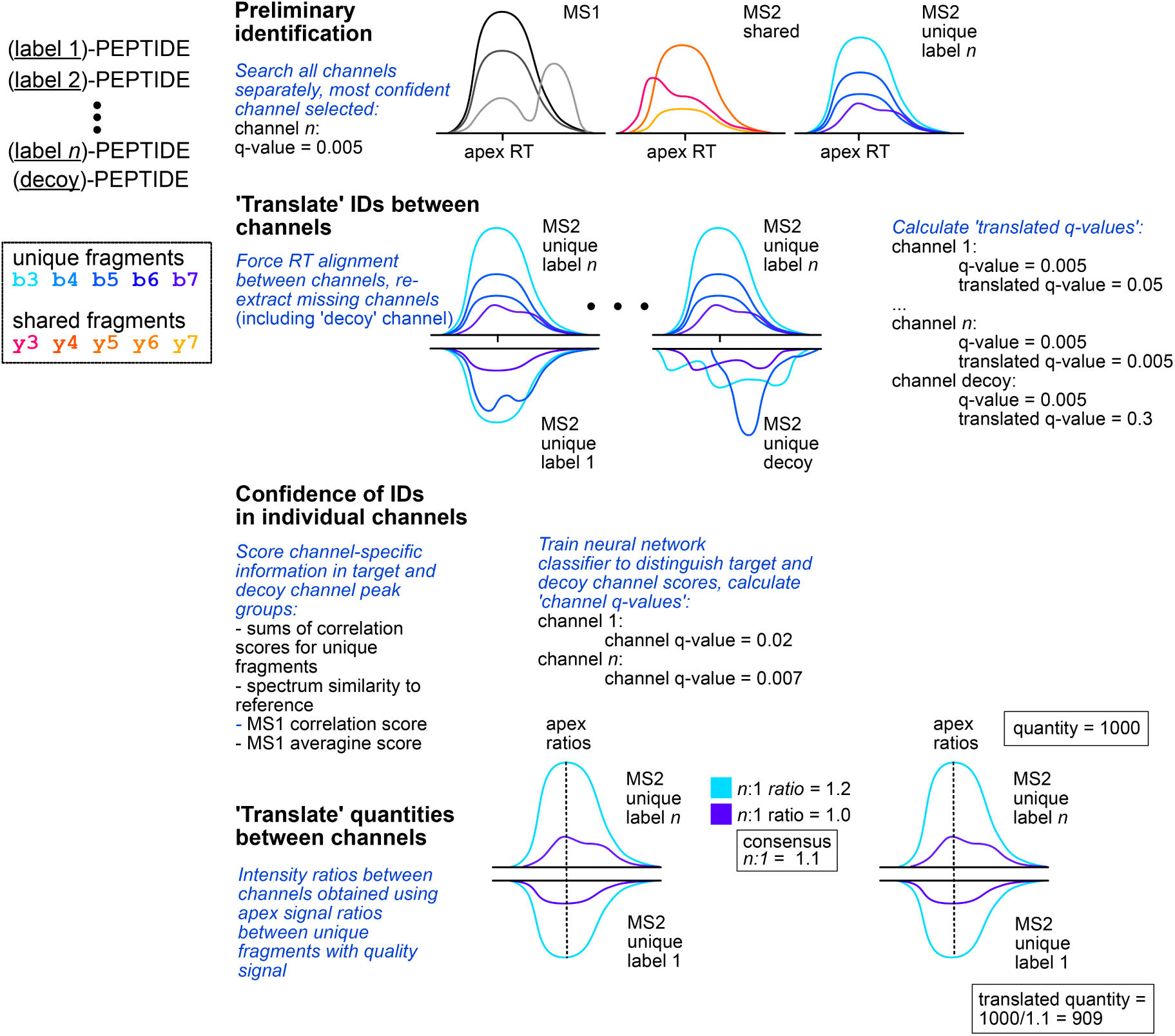
Translated precursor identification and quantification with plexDIA.

**Figure S2.**
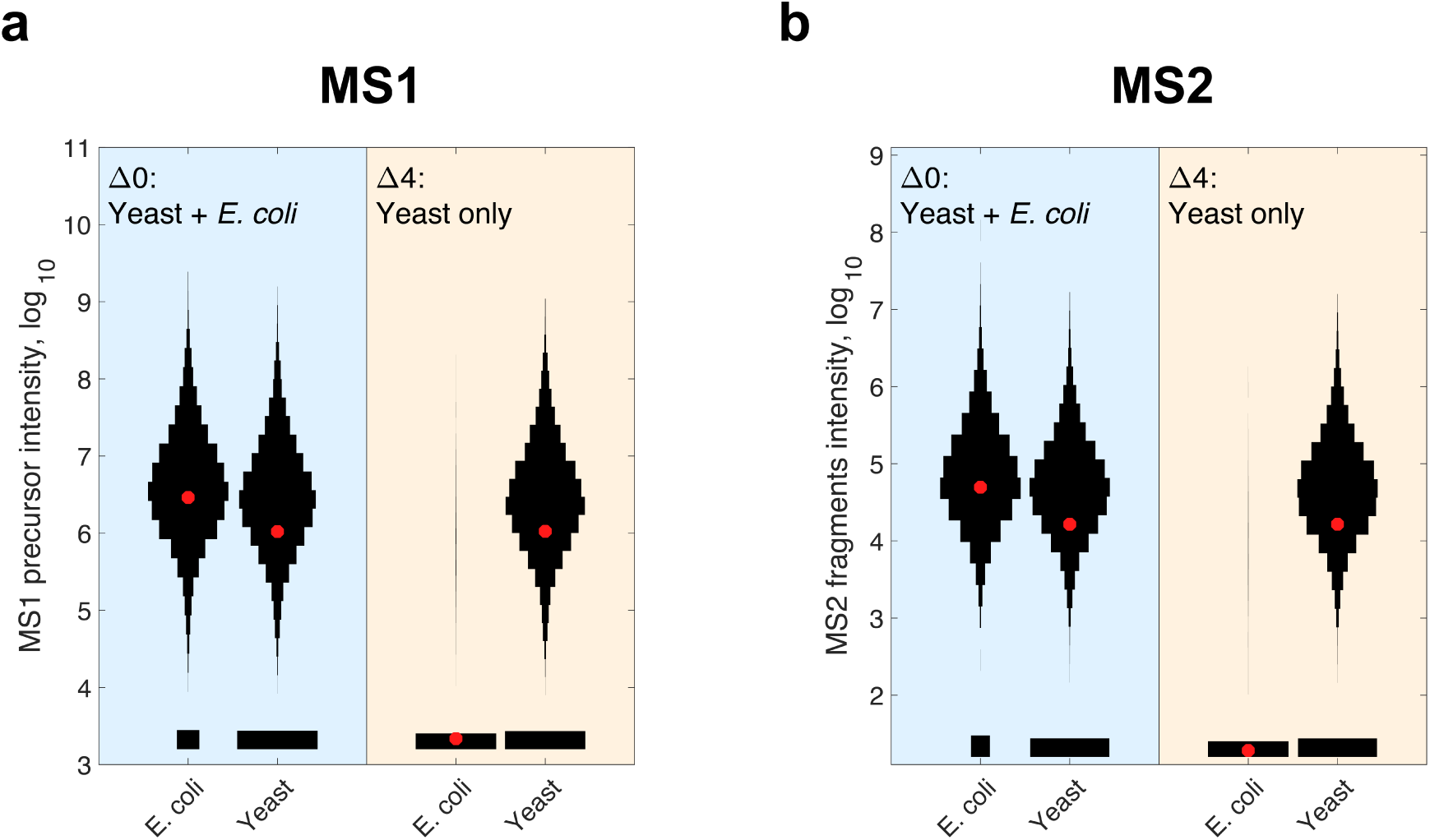
plexDIA analysis of proteins present only in one samples but missing from another. We sought to test identification propagation by plexDIA for the case when proteins are present only in some samples and not in others. To do so, we prepared a standard in which one sample (labeled with mTRAQ Δ0) had both 0.5 *µg E. coli* and 0.5 *µg S. cerevisiae* while another (labeled with mTRAQ Δ4) had only 0.5 *µg S. cerevisiae*. The combined set was analyzed by plexDIA using the V1 method. (**a**) Distributions of raw MS1 precursor intensity for *E. coli* and *S. cerevisiae* precursors at translated channel-q-value *<* 0.01. (**b**) Distributions of raw MS2 quantification of precursors filtered for channel-q-value q-value *<* 0.01. The red asterisks correspond to the means of the distributions.

**Figure S3.**
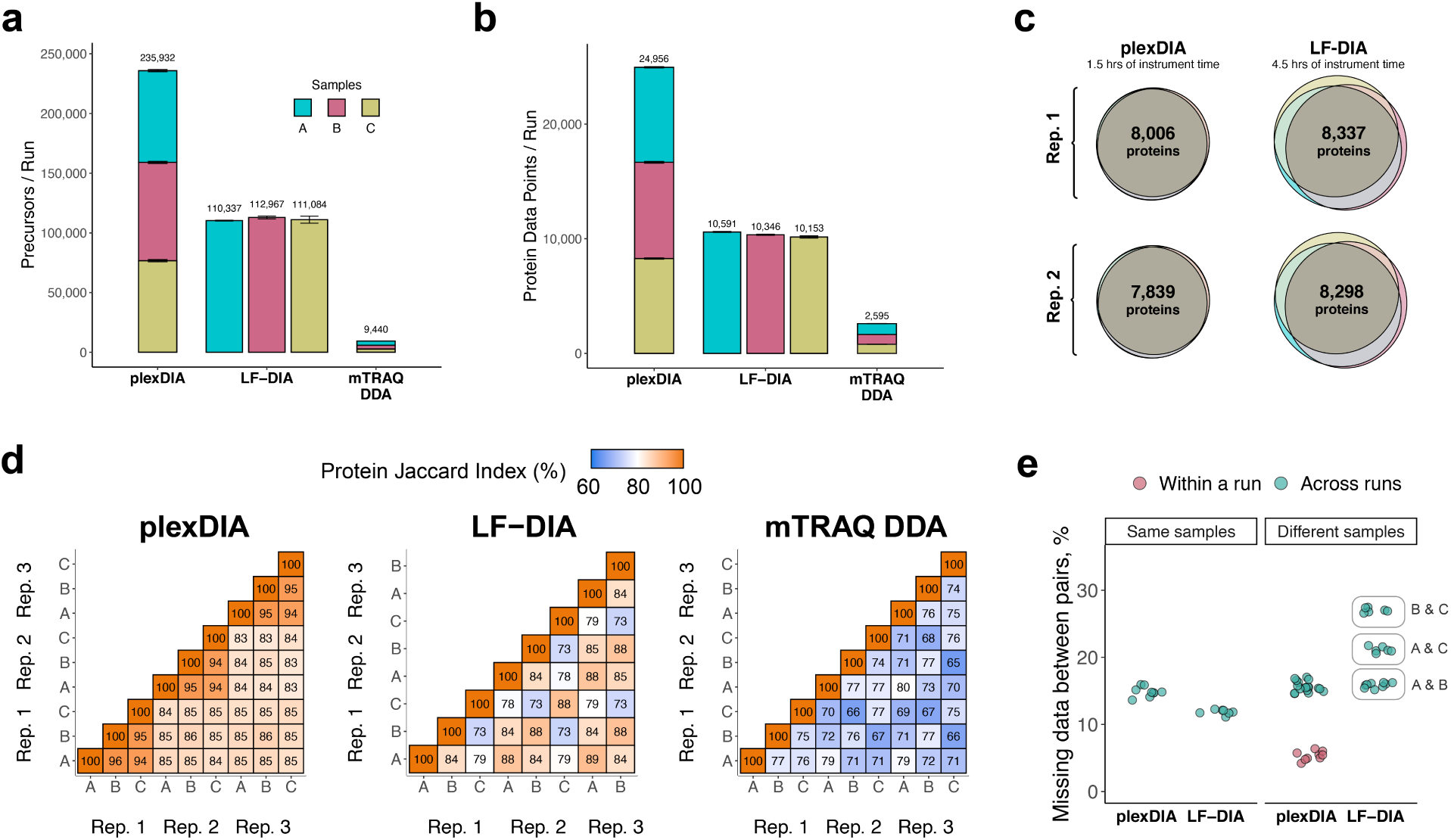
plexDIA proteomic coverage and data completeness for V2. (**a**) Number of distinct precursors identified from 60min active gradient runs for plexDIA, LF-DIA, and shotgun-DDA of mTRAQ at 1 % FDR. The DIA analysis used the V2 method, an MS2-optimized data acquisition cycle shown in Fig. 1. Triplicates of each sample were analyzed (except sample C of LF-DIA, duplicates are analyzed) and the results displayed as mean; error bars correspond to standard error. (**b**) Total number of protein data points for plexDIA, LF-DIA, and mTRAQ DDA at 1 % global protein FDR. (**c**) Venn-diagrams of each replicate for plexDIA and LF-DIA display protein groups quantified across samples A, B, and C. The mean number of proteins groups intersected across samples A, B, and C is 7,923 for plexDIA and 8,318 for LF-DIA(**d**) We compute pairwise Jaccard indices to compare pairwise data completeness between plexDIA, LF-DIA and shotgun DDA for mTRAQ. All data were analyzed using match between runs. (**e**) Distributions of missing data between pairs of runs of either the same sample (i.e., replicate injections) or between different samples

**Figure S4.**
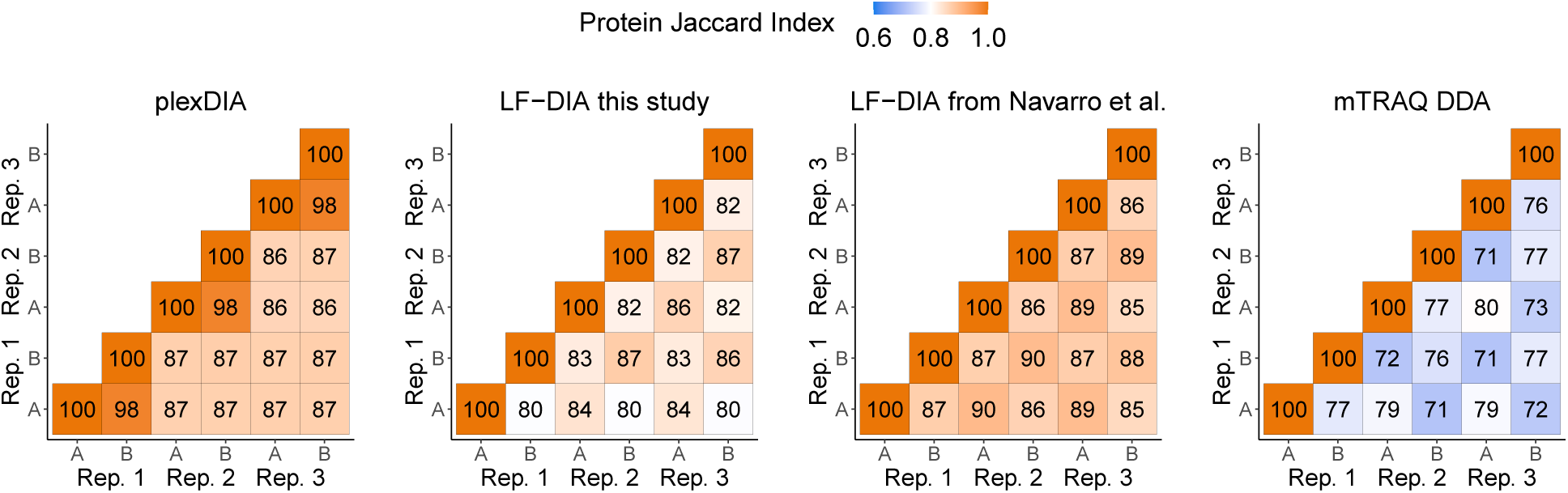
**Comparison of proteomic overlap between our runs to a high quality DIA dataset (Navarro et al.**^23^) DIA runs (including raw data from Navarro et al.) were searched with DIA-NN using match between runs. Results indicate that the data completeness is from LF-DIA in this study is comparable to other high quality LF-DIA datasets.

**Figure S5.**
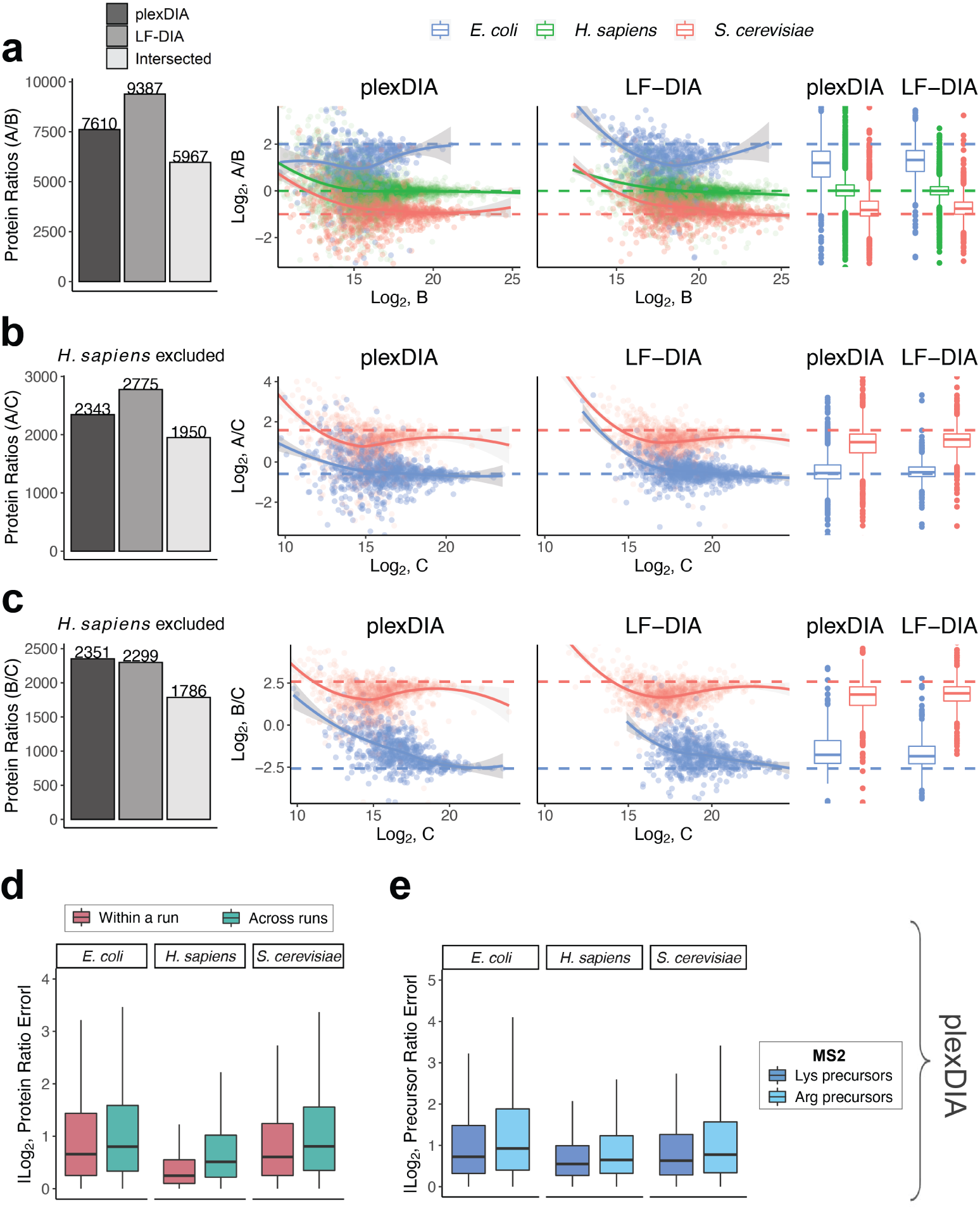
plexDIA quantitative accuracy for MS2-optimized data acquisition (V2) As demonstrated with the MS1-optimized method in Figure 3 of the main text, here we show quantitative accuracy of plexDIA using MS2-optimized data acquisition - specifically, we only show data from the second run of a triplicate set. (**a**) The number of protein groups quantified in both samples A and B is shown with barplots. plexDIA quantified 7,610 PGs, LF-DIA 9,387 PGs, and intersected between plexDIA and LF-DIA was 5,967 PGs. These 5,967 PGs were plotted to compare quantitative accuracy between plexDIA and LF-DIA for in-common protein groups. To improve visibility, the scatter plot x and y axes were set to display data-points between 0.25% and 99.75% range. (**b**) Same as (**a**), but for samples A and C; human proteins were excluded because they compare two different human cell-types. (**c**) Same as (**b**), but for samples B and C. (**d**) Absolute protein ratio errors were calculated for samples A/B, A/C, and B/C and combined to compare ratio errors for samples within a plexDIA run (e.g., run2 A / run2 B), to samples across runs (e.g., run1 A /run2 B) with plexDIA. (e) Absolute precursor ratio errors were calculated for samples A/B, A/C, and B/C and combined to compare MS2-quantified ratio errors for C-terminal lysine precursors and C-terminal arginine precursors.

**Figure S6.**
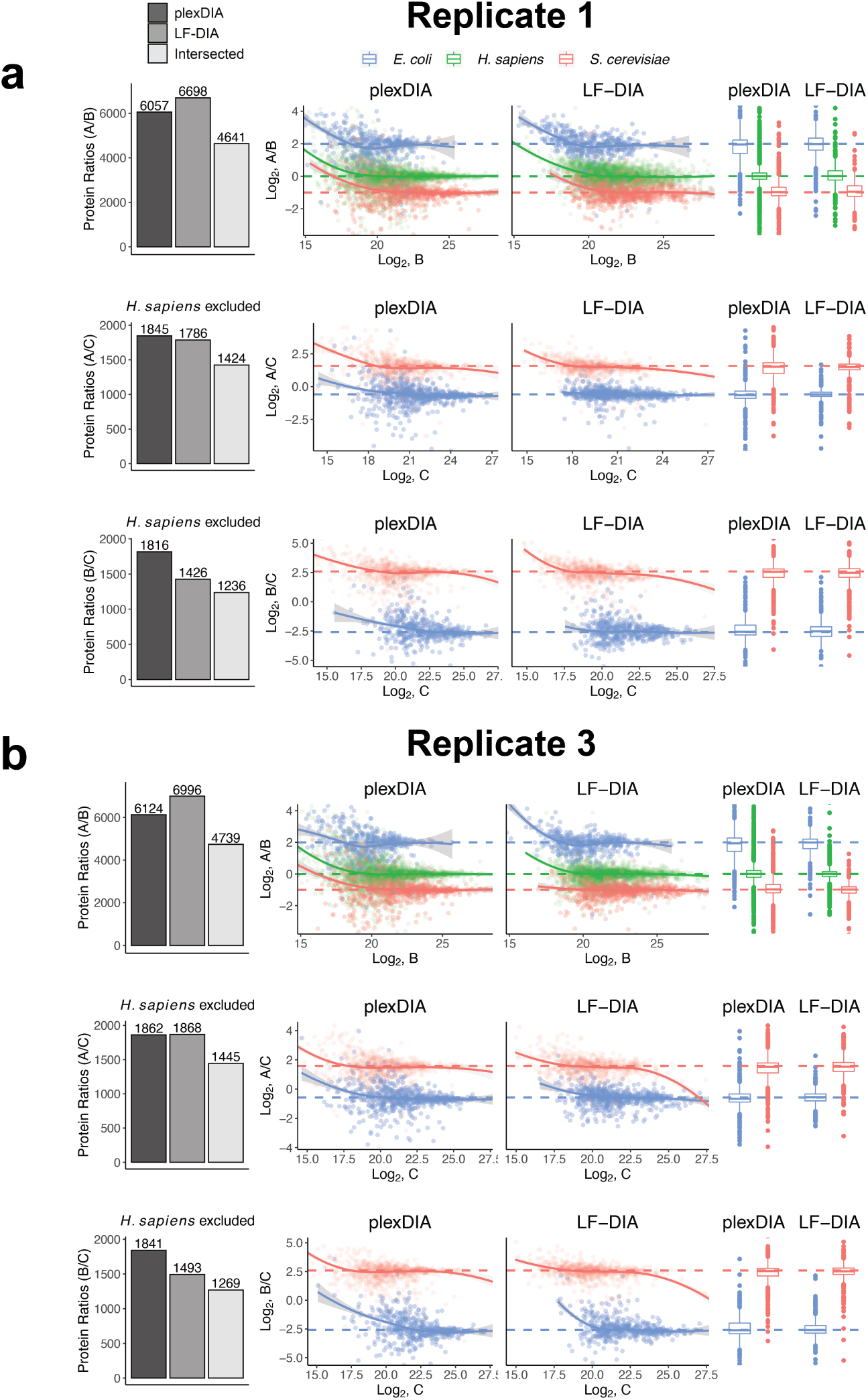
Quantitative accuracy for DIA replicates using V1. Similar to main Figure 3, we display the results from the other replicates. (**a**) Figures are the same as shown in Figure 3 of the main text, with the exception that this shows the first replicate of plexDIA and the first replicate of samples A, B, C for LF-DIA. (**b**) Same as (**a**), but for the third replicate of plexDIA and LF-DIA samples A, B, C.

**Figure S7.**
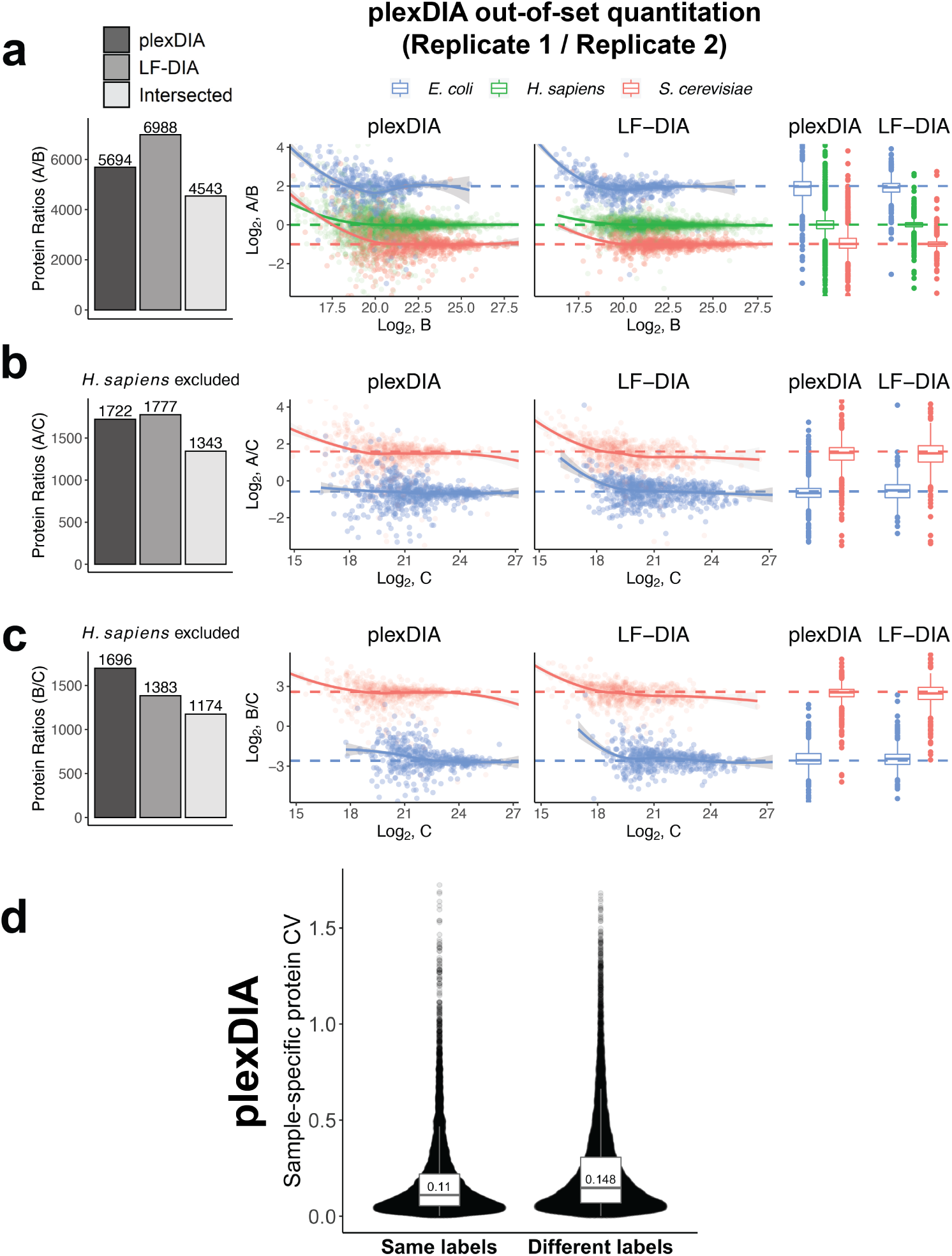
Quantitative accuracy and repeatability across different plexDIA sets and labels. (**a**) Relative protein levels between samples A, B and C estimated from samples analyzed in different plexDIA sets, i.e. out-of-set quantification. The quantitative accuracy between sets (and thus runs) is comparable to the within set accuracy shown in Figure 3. The display is the same as shown in main Figure 3, but the protein ratios are estimated across runs (e.g. run 1 A / run 2 B); LF-DIA is showing protein ratios for the 2nd replicate of samples A,B,C.(**b**) Same as (**a**), but for samples A and C; *H. sapiens* proteins were not analyzed because they are from distinct cell-types. (**c**) Same as (**b**) but for samples B and C. (**d**) Quantitative repeatability of plexDIA across across different labels. Protein CVs were estimated for the same samples labeled with the same label (as in main Fig. 4) or for the same sample labeled with different labels in different runs, e.g. run 1, Δ0, sample A & run 2, Δ4, sample A & run 3, Δ8, sample A. Both distributions contains CV for the same proteins, a set 15,158 sample-specific protein data points per condition (Same Labels or Different Labels). The median CV when using the same label was 0.110 while the label swap had a median CV of 0.148.

**Figure S8.**
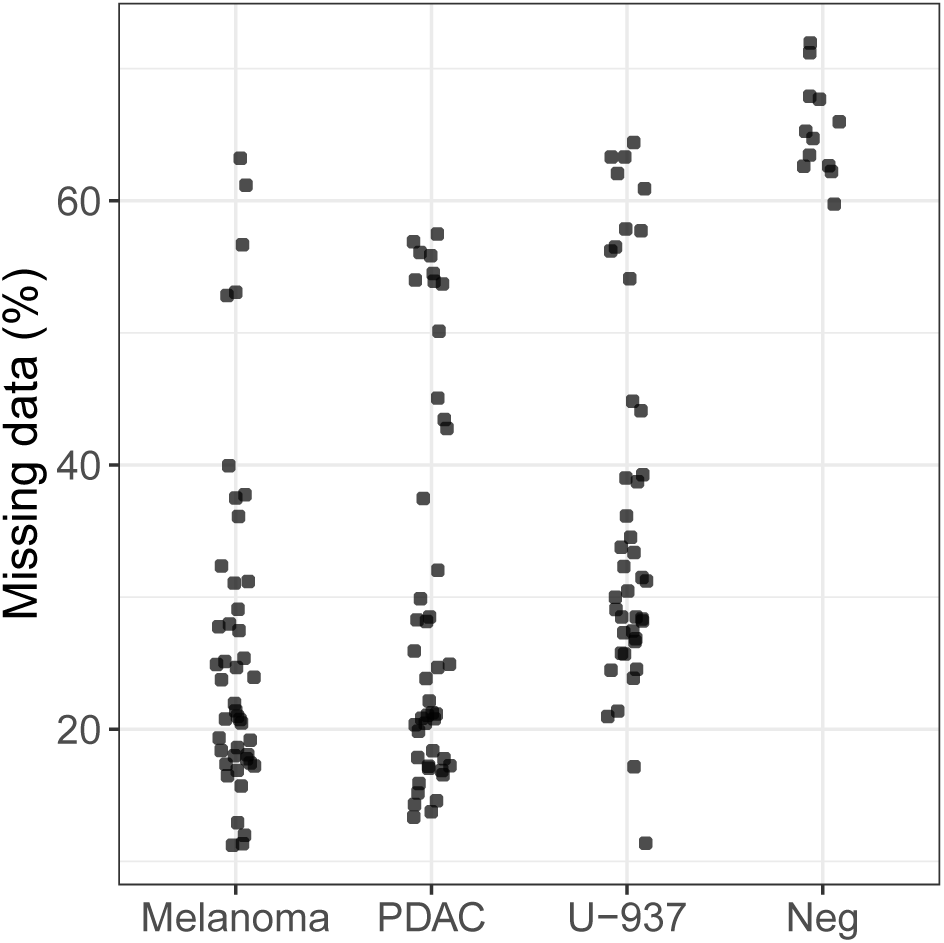
plexDIA missing data in single cells and negative controls. Percent of precursors with no MS1-level quantitation per single cell or negative control. Single cells were required to have *<*60% missing data to be included in downstream analysis.

**Figure S9.**
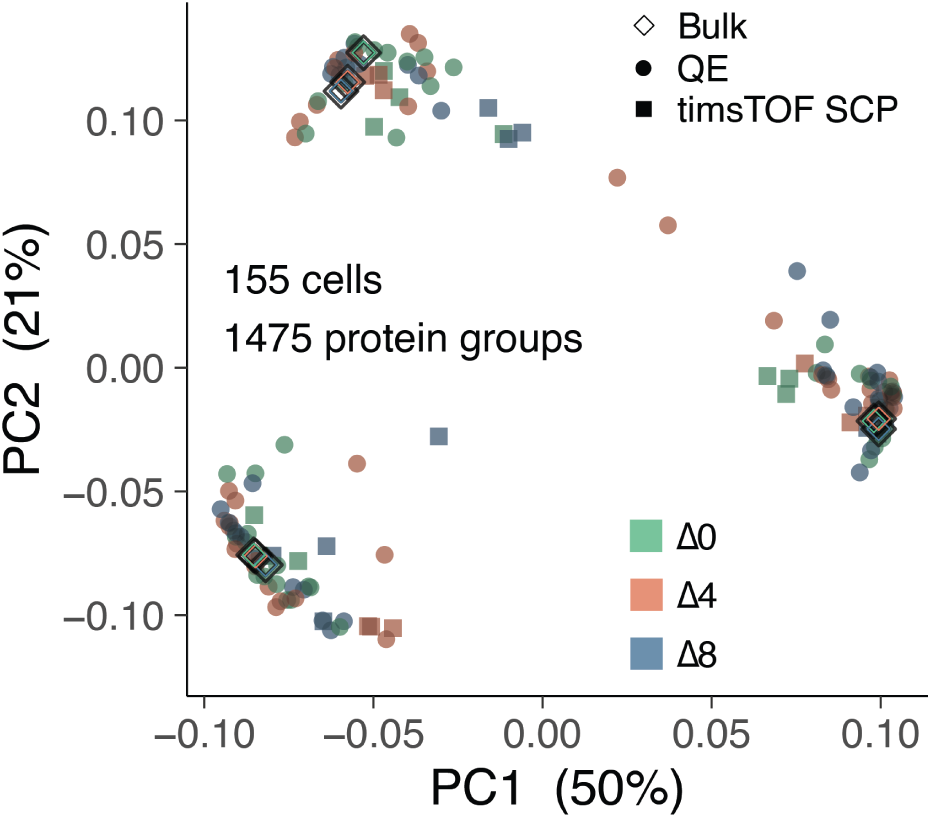
Single-cell PCA colored by mTRAQ label. Rather than colors corresponding to a cell-type as performed in Fig. 6p, here colors correspond to which mTRAQ label was used to tag the single-cells. This is performed to check whether labeling-induced biases affect clustering of single-cells; here there appears to be little to no effect.

**Table S1.** Relative protein abundances for each species per label. Distribution of relative protein abundance of each species across labels. The pooled sample Δ0, Δ4, and Δ8 was used for quantitative benchmarking of plexDIA

| | $\Delta 0$ | $\Delta 4$ | $\Delta 8$ |
| --- | --- | --- | --- |
| *E. coli* | 20% | 5% | 30% |
| *S. cerevisiae* | 15% | 30% | 5% |
| U-937 | 65% | 65% | 0% |
| Jurkat | 0% | 0% | 65% |

